# Development of auxin reporters in oilseed rape (*Brassica napus*)

**DOI:** 10.64898/2026.07.02.736084

**Authors:** Veronika Jedličková, Vendula Pukyšová, Marie Štefková, Matej Zámečník, Marek Sedláček, Hélène S. Robert

## Abstract

Auxin is a key phytohormone that regulates all aspects of plant growth, development, and environmental responses, making the precise analysis of its distribution and signaling essential for understanding plant adaptation and physiological processes. However, despite the agricultural importance of oilseed rape (*Brassica napus*), the lack of robust, species-specific molecular tools limits detailed studies of hormone signaling in this crop. Here, we developed and characterized reporter systems for the sensitive visualization and quantification of auxin distribution and signaling in *B. napus*. The DR5cc auxin signaling reporter and a novel synthetic auxin-responsive reporter, BIP3, assembled from promoter fragments of three oilseed rape *Aux/IAA* genes, were generated to drive GUS expression. In hairy roots, both reporters showed auxin-responsive expression in the root apical meristem that became broader after auxin treatment. In transgenic seedlings, flowers at anthesis, and 12-day-old embryos, DR5cc exhibited a more defined expression pattern than BIP3. To monitor real-time auxin dynamics under abiotic stress, DR5cc fluorescent reporters were employed in hairy roots. Mannitol and NaCl treatments induced a time-dependent increase in fluorescence, peaking at 6–12 h before returning to basal levels after 24 h. Furthermore, dual-reporter assays enabled simultaneous monitoring of auxin and cytokinin signaling, revealing distinct hormone-specific spatial responses in hairy roots. Finally, we established a quantitative DII (qDII) reporter system using degron domains from *B. napus* Aux/IAA proteins, providing a quantitative readout of auxin depletion. Together, these reporter systems enable spatial, temporal, and quantitative analyses of auxin dynamics during development and stress adaptation in oilseed rape.

**Key Message:** Species-specific auxin reporters provide robust tools to visualize and quantify hormone signaling in oilseed rape, advancing studies of development and stress responses.

## Introduction

Plant hormones, or phytohormones, are a diverse group of naturally occurring organic compounds that play crucial roles in regulating plant growth, development, and responses to environmental stimuli. These signaling molecules function at low concentrations and often interact in complex networks to coordinate physiological processes. Insights into hormone interactions and signaling pathways deepen our understanding of plant biology and enhance strategies for crop improvement. Auxin, one of the key phytohormones, regulates numerous developmental processes, including embryogenesis, tropic responses, apical dominance, and cell division, elongation, and differentiation (Di Mambro et al. 2017; Verma et al. 2021; Li et al. 2022). Indole-3-acetic acid (IAA) is the predominant natural auxin, though several other endogenous (e.g., indole-3-butyric acid [IBA], phenylacetic acid) and synthetic auxin analogs (e.g., 2,4-dichlorophenoxyacetic acid [2,4-D], 1-naphthaleneacetic acid [NAA]) also functionally mimic auxin activity (Simon and Petrášek 2011). Auxin signaling is primarily mediated by a pathway that involves TRANSPORT INHIBITOR RESPONSE 1/AUXIN SIGNALING F-BOX (TIR1/AFB) receptor proteins, AUXIN RESPONSE FACTOR (ARF) transcription factors, and Aux/IAA repressor proteins. In the absence of auxin, Aux/IAA proteins inhibit ARF activity. Auxin binding to TIR1 enhances its affinity for Aux/IAAs, promoting their ubiquitination and proteasomal degradation, thereby releasing ARFs to activate target gene expression (Yu et al. 2022). TIR/AFB receptors exhibit adenylate cyclase activity, catalyzing the local production of the second messenger cAMP, which may contribute to transcriptional regulation (Qi et al. 2022; Chen et al. 2025).

Auxin distribution can be visualized directly using immunolocalization, labeled auxins, or biosensors. However, indirect monitoring through transcriptional reporters remains the most widely used approach due to its simplicity and accessibility (Jedličková et al. 2022a). These systems employ natural or synthetic promoters containing auxin-response elements (AuxREs) to drive reporter gene expression, which depends on the presence of auxin signaling components (e.g., receptors, ARFs). As such, these reporters reflect auxin-responsive transcriptional activity rather than auxin concentration *per se*. Although these systems were initially established in *Arabidopsis* (Isoda et al. 2021), they have been increasingly adopted in other plant species, including maize (Robil and McSteen 2023), rice (Yang et al. 2017), and tomato (Yang et al. 2023).

Ulmasov et al. (1995) showed that AuxREs typically contain the core hexamer 5′-TGTCTC-3′, which mediates transcriptional responses to auxin. Building on this sequence, the DR5 reporter was created as a synthetic promoter composed of multiple tandem repeats of an 11-bp sequence (5′-CCTTTTGTCTC-3′) fused to a minimal CaMV 35S promoter (Ulmasov et al. 1997). To enhance signal strength and spatial coverage, the DR5rev variant was developed by arranging nine AuxRE repeats in reverse orientation (Friml et al. 2002). Subsequent structural studies revealed that ARF1 and ARF5 proteins preferentially bind a related motif, 5′-TGTCGG-3′ (Boer et al. 2014). Incorporating this higher-affinity sequence, DR5v2 replaced the TGTCTC repeats with TGTCGG, yielding a promoter with greater ARF binding and improved sensitivity to auxin (Liao et al. 2015).

In plant promoters, AuxREs can occur as pairs of TGTCNN motifs arranged as direct, inverted, or everted repeats, with varying spacing between the sites. In *Arabidopsis thaliana*, the direct repeat with a five-base-pair spacer is most strongly associated with auxin-responsive gene activation, highlighting its role as a highly effective regulatory architecture for auxin-dependent transcription (Freire-Rios et al. 2020). AuxRE variants (TGTCNN) were screened across angiosperms, with TGTCCC, TGTCGG, TGTCGA, and TGTCTC identified as the most highly conserved (Lieberman-Lazarovich et al. 2019). Gene ontology analysis linked regulatory regions containing the TGTCGG motif to cell wall biosynthesis genes and those with the TGTCCC motif to auxin-responsive genes. To test functional relevance, DR5 promoters incorporating individual variants (TGTCCC, TGTCTC, TGTCAC, or the nonconserved TGTCGC) were engineered and reporter activity was assessed in Arabidopsis. All constructs showed dose-dependent auxin responsiveness, but the response strength varied markedly, being highest for TGTCCC (Lieberman-Lazarovich et al. 2019).

In addition to synthetic promoters, promoter fragments from auxin-responsive genes have also been employed as auxin reporters. Examples include promoter regions of *GRETCHEN HAGEN 3* (*GH3*) and *SMALL AUXIN-UP RNA 10A* (*SAUR10A*), which have been used to drive the expression of β-glucuronidase (GUS) or fluorescent proteins (Hagen et al. 1991; Li et al. 1991; Pacios-Bras et al. 2003). The pIAAmotif synthetic auxin reporter was designed with native TGTCCC-containing promoter fragments from *Arabidopsis thaliana IAA1/IAA2* and *Populus trichocarpa IAA1* positioned upstream of a minimal CaMV 35S promoter. Its expression pattern was tested in Arabidopsis and tomato, revealing an overlapping yet broader expression domain compared with the classical DR5 reporter in both roots and shoots (Lieberman-Lazarovich et al. 2019).

Distinct from promoter-based reporters, degron-based systems provide a direct and rapid means of monitoring auxin input by exploiting the auxin-dependent degradation of domain II (DII) in Aux/IAA proteins. In the DII-VENUS system, the DII of IAA28, containing a degron motif required for auxin-induced degradation, is fused to the fast-maturing yellow fluorescent protein VENUS. Auxin triggers TIR1/AFB-mediated ubiquitin–proteasome degradation of the fusion, producing a fluorescence signal inversely correlated with cellular auxin levels (Vernoux et al. 2011; Brunoud et al. 2012). A mutated, auxin-insensitive mDII variant was designed to serve as a stable control. The ratiometric R2D2 system further improves quantification by co-expressing DII-VENUS and mDII-tdTomato, each driven by its own RPS5A promoter within a single construct (Liao et al. 2015). The quantitative DII-VENUS (qDII) variant was developed to measure auxin distribution in the Arabidopsis shoot apical meristem (Galvan-Ampudia et al. 2020). In this system, a single RPS5A promoter drives the expression of both the auxin-sensitive DII-VENUS and TagBFP, a stable blue fluorescent protein used as a reference for normalization, with the two proteins separated by a 2A peptide.

The degron motif (VGWPP[V/I][R/G]XXR) of the DII domain interacts with TIR1/AFB receptor in an auxin-dependent manner. Variations in the degron sequence directly affect Aux/IAA stability and auxin sensitivity. Regions flanking the degron, including the degron tail and proximal motifs such as the Lys-Arg (KR) dipeptide, further modulate auxin-dependent degradation dynamics (Dreher et al. 2006; Moss et al. 2015; Winkler et al. 2017; Figueiredo et al. 2022; Figueiredo and Strader 2022).

In this study, we established a reporter system that can be used as a sensitive and robust tool for visualization and quantification of auxin levels and response distribution in oilseed rape, a globally important oilseed crop.

## Methods

### Plant material

Seeds of *B. napus* cultivar DH12075 used for transformation experiments were surface-sterilized with 20% bleach and cultivated *in vitro* on half-strength Murashige and Skoog (MS) medium (Duchefa) supplemented with 8 g/L plant agar (Duchefa) and 5 g/L sucrose. Growth conditions were set to 21°C with a 16-h photoperiod in a controlled environment chamber. Plants grown in soil were cultivated in a greenhouse under similar temperature and light regimes.

### Reporters construction

For DR5cc reporter construction, a fragment containing nine tandem AuxRE motifs (TGTCCC) separated by 5-bp spacers, as naturally occurring in the promoters of *IAA* genes in *B. napus* and *A. thaliana*, was synthesized together with a minimal CaMV 35S promoter (Invitrogen). The fragment was cloned into the Addgene plasmid pICH41233 or pICH41295 (Weber et al. 2011) to generate reporter constructs with and without the Ω UTR, respectively. The Addgene plasmids used for L1 and final L2 assembly (Engler et al. 2014) of the GUS reporter and fluorescent variants are detailed in Supplementary Fig. S1. The final plasmids used for plant transformation also contained a gene conferring kanamycin (GUS and Venus variants) (Li et al. 2013) or hygromycin (mCherry variant) (Gantner et al. 2018) resistance.

For BIP3 reporter construction, promoter fragments of *BnaIAA* genes were selected from multiple homoeologous copies present in the genome. For *BnaIAA1* and *BnaIAA4*, the A- and C-subgenome copies (*BnaA08g30190D* and *BnaC08g09640D* for *IAA1*, *BnaA09g16590D* and *BnaCnng63870D* for *IAA2*) were identical within the selected region (Supplementary Fig. S2). For *IAA2*, six copies were identified in the *B. napus* genome (Li et al. 2017). The copy selected for reporter construction (*BnaA01g24190D*, identical to *BnaC01g31190D* in this region) was chosen based on the presence of TGTCCC motifs (Supplementary Fig. S2) and expression data from the Brassica Expression Database (Chao et al. 2020). A composite fragment consisting of three 51–52 bp promoter elements, together with a minimal CaMV 35S promoter, was synthesized (Invitrogen). The vectors were constructed using the MoClo protocol, employing the Addgene plasmids specified in Supplementary Fig. S1.

The DII domains of *IAA2* and *IAA17* (*BnaA01g24190D* and *BnaA08g27770D*, respectively), selected for qDII reporter construction (Supplementary Fig. S3), were synthesized (Eurofins) and cloned into the Addgene plasmid pAGM1276 (Weber et al. 2011). The DII domain was cloned in frame by MoClo with a synthesized Venus–NLS–P2A–TagBFP–NLS fragment (Eurofins). The construct contained the promoter and 5′UTR of CaMV 35S, and the Atug7 terminator (Supplementary Fig. S1) (Engler et al. 2014). The final plasmids used for plant transformation contained a gene conferring kanamycin resistance (Li et al. 2013).

The sequences of all plasmids generated in this study have been deposited in the Zenodo repository (https://doi.org/10.5281/zenodo.21030825).

### Plant transformation and regeneration

Hairy root induction was performed using the Ti plasmid–less *Agrobacterium tumefaciens* strain C58C1 carrying the hairy root–inducing plasmid pRiA4b (Petit et al. 1983), and *A. rhizogenes* strain K599. Reporter constructs were introduced into *Agrobacterium* cells by electroporation. Transformation of *B. napus* hypocotyls was carried out using an injection-based method, as described in Jedličková et al. (2023). Independent hairy root lines were cultured on solid MS medium supplemented with Gamborg B5 vitamins (Duchefa) and 30 g/L sucrose. The medium was further supplemented with cefotaxime (200 mg/L) to suppress bacterial growth and kanamycin (20 mg/L) or hygromycin (15 mg/L) for selection of reporter constructs. Cultures were maintained at 22°C in the dark. Regeneration of selected hairy root lines was performed according to an optimized protocol using 5 mg/L 6-benzylaminopurine (BAP) and 8 mg/L 1-naphthaleneacetic acid (NAA) at 21°C under a 16-h photoperiod (Jedličková et al. 2023). The regenerated plants (T0) were backcrossed with wild-type DH12075, and the resulting T1 progeny were screened by PCR using DNA extracted from leaves via the conventional cetyltrimethylammonium bromide (CTAB) method (Allen et al. 2006), with specific primers targeting *GUS*, *rolA* and *aux1* genes (encoded by TL and TR of *Ri* plasmid, respectively; Supplementary Table S1).

### Transgene copy number assessment

A 0.6 kb fragment of the reference gene *ACC* was amplified from genomic DNA of DH12075. Two distinct copies of the gene were obtained by cloning. Sequence alignment revealed 82% similarity between the two copies (Supplementary Fig. S4). One copy originated from the A subgenome (99.51% similarity to *ACC* in *B. rapa*), whereas the other was derived from the C subgenome (98.18% similarity to *ACC* in *B. oleracea*). The cloned fragments of the *ACC*-A and *ACC*-C copies were subsequently used to optimize qPCR conditions to specifically amplify the *ACC*-A copy. The qPCR program consisted of an initial step at 50 °C for 2 min, followed by 90 °C for 3 min, and 40 cycles of 95 °C for 15 s, 60 °C for 10 s, and 72 °C for 18 s. Reactions were performed in a total volume of 10 μL, containing 5 μL of 2× PrimeTime Gene Expression Master Mix (IDT), 0.25 μL each of forward and reverse primers (10 μM), 0.06 μL of the probe (10 μM), 2 μL of diluted DNA, and 1.94 μL of nuclease-free water. Amplification was carried out using an Applied Biosystems QuantStudio 12K Flex system. Primers and probes are specified in Supplementary Table S1. Standard curves for *ACC*-A and *uidA* were generated using five successive dilutions of oilseed rape genomic hairy root DNA (625, 125, 25, 5, and 1 ng per reaction). The slope and intercept of these standard curves were used to calculate transgene copy number in each hairy root line according to the formula described by Weng et al. (2004). Analyses were performed using three biological replicates, each comprising three technical replicates.

### Histochemical GUS staining

Samples from regenerated plants (4-day-old seedlings, flowers, 12-day-old dissected embryos) were immersed in GUS staining solution (0.5 M sodium phosphate buffer; pH 7, 50 mM potassium ferricyanide (III) [Sigma], 0.2% Triton X-100 [Serva], 20 mg/ml X-Gluc [Thermo Fisher Scientific] in DMSO), incubated in vacuum for 15 min, and stained for 16 h at 37 °C. After staining, samples were washed in phosphate buffer and submerged in 70 % ethanol to remove chlorophyll. Embryos were cleared with chloralhydrate solution (8/3/1 w/v/v chloralhydrate [Lach-Ner], H_2_O, Glycerol [Penta]). Transgenic hairy roots grown on plates were excised (1-cm segments), incubated in GUS staining solution for 2 h at 37 °C and directly imaged.

### Hormonal and abiotic stress treatment

Root tips of transformed *B. napus* hairy roots were collected 10 days after subculture and immersed in liquid MS-B5 medium supplemented with the respective chemicals or an equal volume of DMSO (mock treatment). To assess the auxin sensitivity of the fluorescent DR5cc reporter, plants were treated with 1 µM NAA or 1 µM IAA, and images were acquired after 6 and 24 h. Cytokinin sensitivity was assessed by treating samples with 5 µM BAP for 2 h. For the combined treatment, root tips were first exposed to 1 µM NAA for 22 h, followed by an additional 2 h treatment with 1 µM NAA and 5 µM BAP. To assess auxin distribution in hairy root tips expressing the qDII reporter, 10 µM NAA was applied for 2 h. The IAA dose response of the qDII reporter was assessed by treating hairy roots with mock or 1, 5, or 10 µM IAA. For stress response analysis, hairy roots were immersed in MS-B5 medium containing 75 mM NaCl (Penta) for salinity treatment and 150 mM D-Mannitol (Sigma) for osmotic stress. MS-B5 medium alone was used for the mock treatment. Samples were incubated at constant temperature of 21°C for 1, 6, 12, and 24 h. To assess TCSv2 cytokinin sensitivity in regenerated roots, 4-old-day seedlings were immersed in MS media containing 10 µM BAP for 2 h. Root tips were then collected and fixed in 4% PFA (Merck) in PBS-T (1X PBS with 0.05% Triton X100 [Serva] pH 7.4) for 1 h under vacuum conditions, followed by three washes of 1 h each in PBS-T (pH 7.4). Samples were transferred to a fresh ClearSee alpha solution (10% (w/v) xylitol [Sigma], 15% (w/v) sodium deoxycholate [TCI], 25% (w/v) urea [Merck], 100 mM sodium sulfite [Sigma] in water). Images were acquired after 5 days of clearing at room temperature.

### Microscopy

Images of GUS-stained samples were acquired using Olympus SZX16 stereomicroscope and Zeiss Axioscope.A1 ZEN histological microscope. Confocal images of hairy root tips were taken using Zeiss LSM 880 confocal microscope equipped with 25x objective. Three laser lines were used in the study: 405 nm for TagBFP (detection at 410-507 nm), 514 nm for VENUS (detection at 519-586 nm), and 561 nm for mCherry (detection at 578-696 nm). Scans were performed at 1x zoom and in 1094*1094 or 2048*2048 pixel resolution. Laser intensity, detector gain, and pinhole size were kept constant within each replicate with pixel time around 1 µs. Maximum intensity projections of confocal z-stacks were acquired at 5.5 µm intervals; the number of slices varied depending on root thickness. Imaging of the qDII hairy roots treated with IAA was performed using Olympus IXplore SpinSR spinning-disc confocal microscope system (Evident) equipped with a Yokogawa CSU-W1 scanner unit (using the 50 µm pinhole disk). Samples were imaged with a 40x objective. TagBFP and VENUS signals were excited using 405 nm and 515 nm diode lasers, respectively. Emitted fluorescence was collected through appropriate emission filter sets onto an sCMOS camera. Z-stacks were acquired with a 1.5 µm step size, and scans were performed in 2034*2034 pixel resolution. Laser power and exposure time were optimized for each reporter line and kept constant across mock and treated samples. The original uncropped and unprocessed images have been deposited in the Zenodo repository (https://doi.org/10.5281/zenodo.21030825).

### Measurements and statistical analysis

Fluorescence intensity was quantified using mean gray value measurements from defined regions of interest (ROI). For the DR5 and TCSv2 reporters, background fluorescence was measured for each image and subtracted prior to further analysis. For the NAA-treated qDII hairy roots, single-plane confocal images of the epidermis were analyzed using ImageJ software (Schneider et al. 2012). Fluorescence was quantified in 15 individual nuclei expressing both stable TagBFP and degradable VENUS. For the IAA dose-response assays, ratiometric analysis was performed on confocal z-stacks using IMARIS 10.2.0. (Bitplane, http://www.bitplane.com/imaris/imaris). Representative images generated using IMARIS are presented. Nuclei were segmented and masked using the stable TagBFP channel within a standardized ROI above the root tip, and mean fluorescence intensities for VENUS and TagBFP were extracted from the masked nuclei. For both pipelines, the VENUS / TagBFP signal ratios of all segmented nuclei within the ROI were averaged. In all assays, an individual hairy root was defined as a biological replicate (n). Resulting values were normalized to the average signal of mock control, and data are presented as relative fluorescence intensity. A total of 3 - 4 independent experiments were performed for each analysis. Depending on the assay, one-way ANOVA followed by Dunnett’s test or a Welch’s t-test were performed to determine differences in relative fluorescence intensities in the roots under mock and treated conditions.

## Results

### A DR5cc auxin reporter for oilseed rape

Building on the previously designed synthetic DR5 promoter (Ulmasov et al. 1997), we developed a variant of this reporter for oilseed rape. In place of the original core AuxRE motif TGTCTC, we introduced the TGTCCC motif, which has been shown to confer high auxin responsiveness in Arabidopsis (Lieberman-Lazarovich et al. 2019), and designated the construct as DR5cc. As in the original DR5 construct, nine repeats of the core motif are separated by 5-bp spacers. To reduce repetitive sequence composition, however, we supplemented the previously used CCTTT spacer (Ulmasov et al. 1997; Lieberman-Lazarovich et al. 2019) with additional 5-bp sequences naturally occurring in *Aux/IAA* promoters of oilseed rape and Arabidopsis. For example, the ATCTT sequence, found between two core motifs in the *BnaIAA1* promoter, was incorporated between the first two TGTCCC motifs in DR5cc (Fig. 1A). The DR5cc sequence was fused to the minimal CaMV 35S promoter to drive expression of β-glucuronidase (GUS). Two constructs were generated, differing in the presence or absence of the 5′-untranslated region (5′-UTR) of tobacco mosaic virus (TMV) RNA (Ω), and designated as DR5cc-GUS and DR5cc-Ω-GUS (Fig. 1). In addition, we included a negative control containing only the minimal CaMV 35S promoter and a positive control in which GUS expression was driven by the full CaMV 35S promoter.

**Fig. 1.**
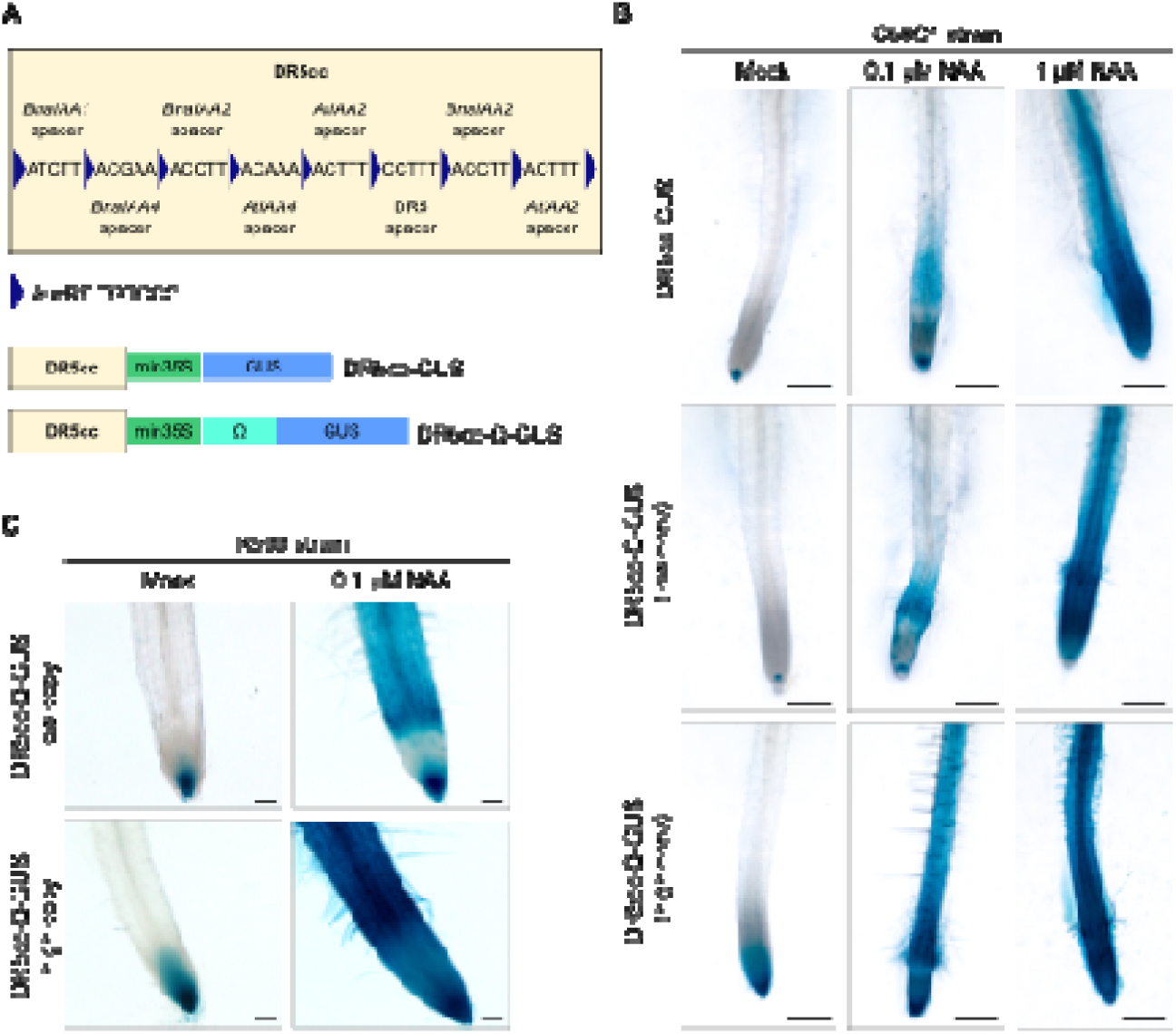
DR5cc reporter in oilseed rape. (A) Schematic representation of DR5cc constructs transformed into *Brassica napus* DH12075. (B) Hairy root lines induced by the *Agrobacterium* strain C58C1 carrying DR5cc-GUS and DR5cc-Ω-GUS constructs, illustrating endogenous auxin signaling (without induction) and responses following induction by exogenous auxin treatment (NAA at 0.1 µM and 1 µM for 12 h). For the DR5cc-Ω-GUS constructs, both low-copy (1 copy) and high-copy (9 copies) lines are shown. For the DR5cc-GUS construct, a line containing 3 copies is presented. Scale bars represent 500 µm. (C) Hairy root lines induced by the *Agrobacterium* strain K599 carrying the DR5cc-Ω-GUS construct, analyzed under non-induced conditions and after induction with 0.1 µM NAA. The low-copy line contains 1 copy, whereas the high-copy line contains 6 copies of the transgene. Scale bars represent 200 µm

*B. napus* DH12075 was transformed using C58C1 *Agrobacterium* strain harboring *pRiA4* plasmid and one of the binary vectors. Independent hairy root lines were subsequently screened for GUS activity. In the hairy root lines under study, the transgene copy number was determined using quantitative PCR (qPCR). Following previous studies (Hernández et al. 2001; Long et al. 2022), the *acetyl-CoA carboxylase* (*ACC*) gene was selected as an endogenous control. Primers and a probe for qPCR were designed to selectively amplify the A-subgenome copy (*ACC-A*). The transgene (*uidA* gene commonly referred to as the GUS gene) copy number in hairy roots was then determined following the approach described by Weng et al. (2004).

Hairy root lines carrying the DR5cc-GUS and DR5cc-Ω-GUS constructs exhibited similar pattern of auxin signaling (Fig. 1B, Supplementary Fig. S5). In the absence of exogenous auxin, GUS activity was detected in the root apical meristem (RAM). Following auxin treatment (0.1 µM and 1 µM NAA for 12 h), the GUS signal became widely distributed. As expected, hairy roots carrying the minimal CaMV 35S promoter did not show any GUS activity either before or after auxin treatment, whereas the positive controls containing the full CaMV 35S promoter exhibited strong signal throughout the root (Supplementary Fig. S5).

In addition, *B. napus* DH12075 was transformed using the K599 *Agrobacterium* strain harboring the DR5cc-Ω-GUS construct (Fig. 1C). The transformation efficiency using the injection-based method was 92.6 ± 2.6%, which was similar to that obtained with the C58C1 strain (96.7 ± 2.9%) (Jedličková et al. 2022b). The difference between the hairy root lines transformed with K599 and those transformed with C58C1 became evident during growth in tissue culture. Whereas the C58C1 lines exhibited vigorous growth, the K599 lines developed slowly, with some lines performing very poorly or dying on the plates (Supplementary Fig. S6).

No major differences in GUS localization were observed between DR5cc-Ω-GUS lines generated with the C58C1 and K599 strains. However, the transgene copy number appears to influence the intensity and diffusion of the GUS expression pattern in lines produced by both *Agrobacterium* strains, with lines containing multiple GUS copies showing stronger GUS intensity than those with fewer transgene copies (Fig. 1B and 1C).

### A new synthetic auxin reporter specific for oilseed rape

To better mimic the context of a native promoter, we designed an auxin reporter comprising three fragments, each 51 – 52 bp long, derived from *B. napus* promoters *BnaIAA1*, *BnaIAA2*, and *BnaIAA4*. Each fragment contains the native TGTCCC AuxRE motif (Fig. 2A). As oilseed rape is an allotetraploid species, it contains multiple copies of each gene derived from the A and C subgenomes. The copies chosen for the design of the reporter were identified based on the presence of TGTCCC motifs in their sequences (Supplementary Fig. S2) and supporting expression data (Chao et al. 2020). Interestingly, the *BnaIAA4* promoter contains a third TGTCCC motif arranged as an inverted repeat downstream of the second TGTCCC motif, separated by 9 bp (Fig. 2A). The construct was designated as BIP3 (*Brassica IAA* promoter – 3x). The synthetic promoter was fused to a minimal CaMV 35S promoter driving GUS expression, and two variants were generated, including the Ω 5’ UTR (BIP3-Ω-GUS) or not (BIP3-GUS).

**Fig. 2.**
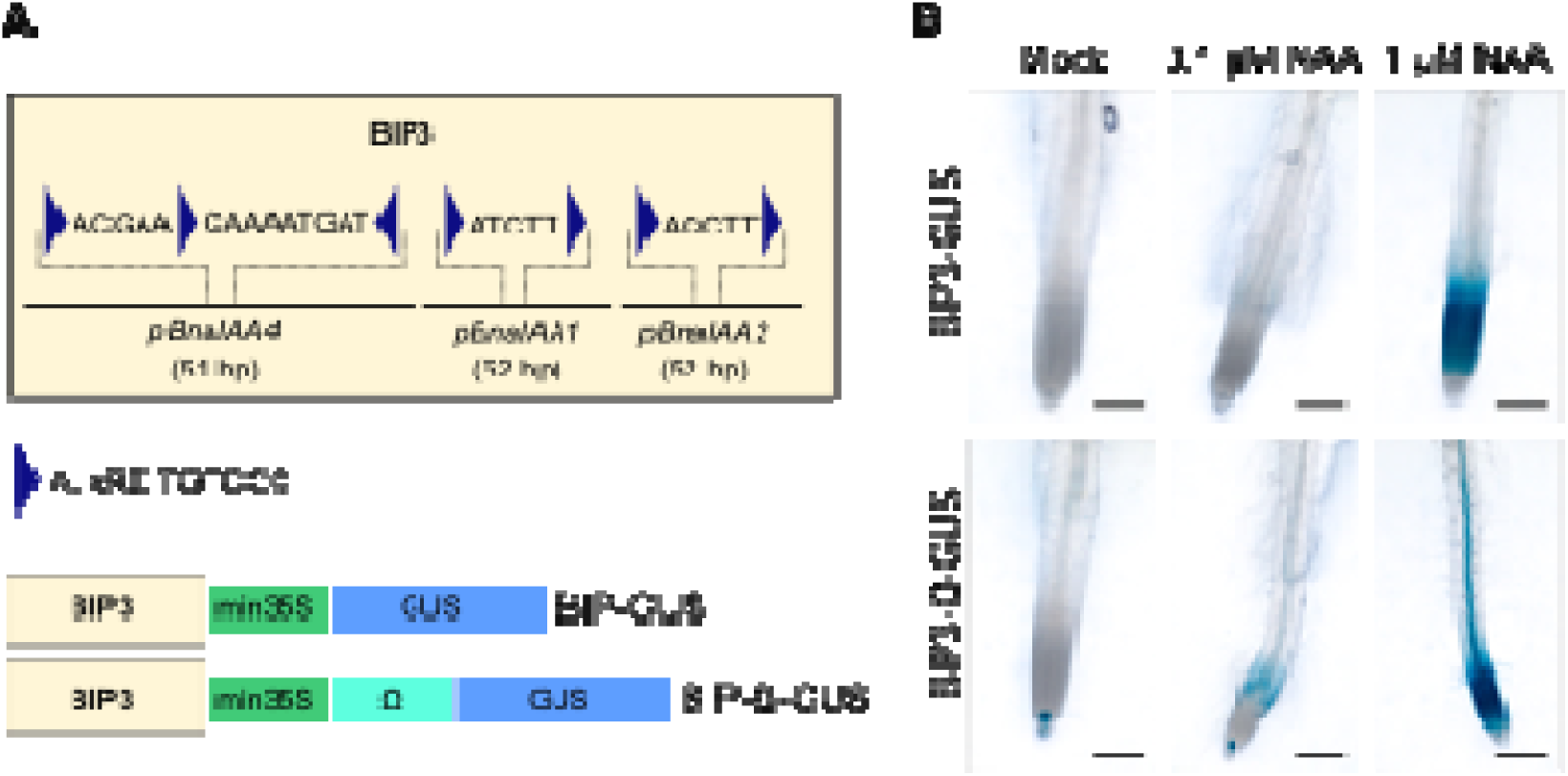
BIP3 reporter in oilseed rape. (A) Schematic representation of the BIP3 auxin reporter composed of fragments derived from three *BnaIAA* promoters containing AuxREs as either direct or inverted repeat. (B) Detection of auxin signalling in hairy roots transformed with *Agrobacterium* C58C1 carrying the BIP3-GUS or BIP3-Ω-GUS constructs following induction with 0.1 µM or 1 µM NAA for 12 h, or in the absence of induction (Mock). Both lines contain 3 copies of the transgene. Scale bars represent 500 µm

In the BIP3 reporter lines, endogenous auxin activity was weakly reported in the RAM. Application of 0.1 µM and 1 µM NAA for 12 h broadened the GUS signal, showing a clear concentration-dependent gradient, with the 1 µM treatment producing a stronger and more extensive response (Fig. 2B). GUS staining patterns were consistent across independent hairy root lines (Supplementary Fig. S5).

### Auxin signaling in regenerated plants

Following analysis in hairy roots, we regenerated selected hairy root lines carrying the DR5cc-GUS, DR5cc-Ω-GUS, BIP3-GUS, and BIP3-Ω-GUS constructs. The T0 regenerants derived from transformation with the C58C1 strain exhibited the typical *Ri* phenotype, often characterized by a dwarfed stature and curly leaves (Christey 2001; Jedličková et al. 2022b). These T0 regenerants were backcrossed to the DH12075 wild type (WT). The resulting backcrossed T1 plants were genotyped for the presence of T-DNAs derived from the binary vector (GUS transgene) and from the *Ri* plasmid. In the *pRiA4* plasmid, the T-DNA is divided into two separate regions, the left (TL) and the right (TR), which were analyzed independently. Of the 244 plants screened, derived from six independent transgenic lines carrying either the DR5cc or BIP3 constructs, the majority (79.5%) contained all three T-DNAs (+GUS, +TL, +TR). No T-DNA was detected in 10.7% of the plants (−GUS, −TL, −TR), while 9.8% lacked the TR region (+GUS, +TL, −TR). Notably, none of the analyzed plants exhibited a genotype containing the GUS transgene while lacking both *Ri* T-DNA regions (+GUS, −TL, −TR).

The expression pattern of the auxin signaling-based reporters was examined across different tissues in T1 plants, including 4-day-old seedlings, root tips, flowers at anthesis, and 12-day-old embryos (Fig. 3). As expected, the positive control harboring the full CaMV 35S promoter showed ubiquitous GUS expression, whereas no signal was detected in the WT plants (Fig. 3A).

**Fig. 3.**
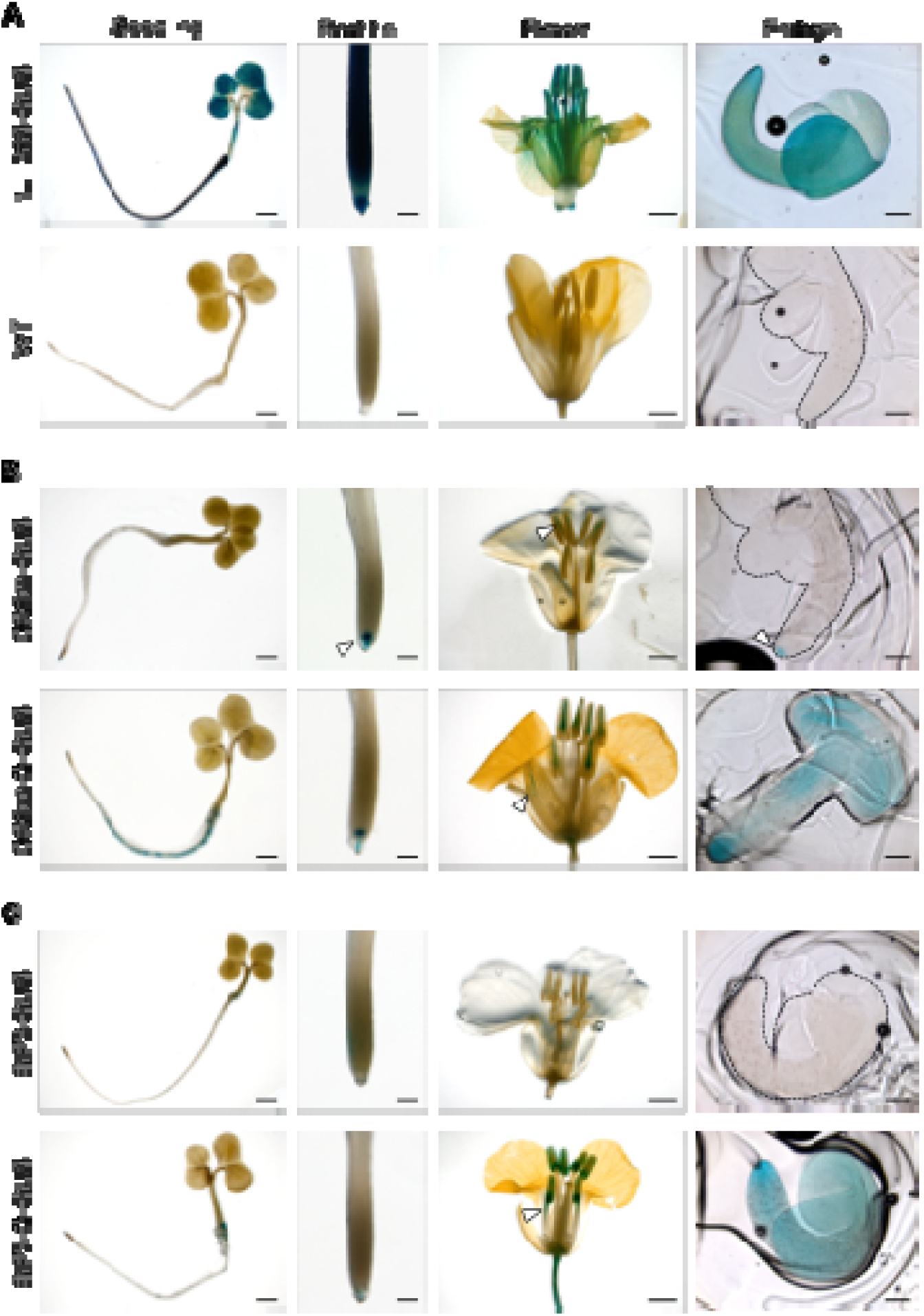
Histochemical GUS staining of different tissues of *B. napus* transgenic plants. (A) positive and negative controls. (B) DR5cc-GUS (+/- Ω). (C) BIP3-GUS (+/- Ω). Scale bars represent 2 mm (seedlings, flowers), 200 µm (root tips), and 500 µm (embryos). Arrowheads indicate the quiescent centre and columella cells in the root tip, the anther connective tissue in the flower, and the embryonic root pole in the embryo for DR5cc-GUS; the sepal vasculature in the flower for DR5cc-Ω-GUS; and the anther filament in the flower for BIP3-Ω-GUS

DR5cc-GUS reporters exhibited restricted expression in root tips, confined to the quiescent centre (QC) and columella cells (Fig. 3B). In flowers, the signal was detected in the anther connective tissue, while in seeds, it was localized to the embryonic root pole. In contrast, DR5cc-Ω-GUS plants displayed broader and more stable expression. In seedlings, staining extended beyond the root apex to vasculature, lateral root primordia, and the root-hypocotyl junction. In flowers, signal was detected in sepal vasculature and within the anther connective tissue. In seeds, embryos showed widespread strong staining throughout the cotyledons and root.

BIP3-GUS exhibited weak, diffuse staining, with low signal in the root tip, weak staining at the hypocotyl-root junction, faint signal in anther connective tissue, and no detectable staining in embryos (Fig. 3C). Addition of the 5’ UTR in BIP3-Ω-GUS enhanced reporter sensitivity, resulting in more defined staining in columella, stronger signal in vasculature near the root-hypocotyl junction, and expression in pedicel, anther filaments, connective tissue, and anther lobe. In embryos, broad staining was observed, resembling that of a strong auxin reporter. GUS staining patterns were consistent across independent lines harboring the same construct (Supplementary Fig. S7).

Reporter functionality in transgenic plants was further confirmed by treating roots for 3 h with NAA or IAA at concentrations of 0.1 µM and 1 µM, resulting in enhanced GUS staining, and by locally applying 50 µM NAA to inflorescences, where broader staining was observed after 24 h compared with mock-treated controls (Supplementary Fig. S8 – S10).

### Abiotic stress-induced auxin signaling response in oilseed rape

To enable live imaging of auxin responses, we generated fluorescent reporters by fusing the DR5cc-Ω to sequences encoding the VENUS and mCherry proteins, and transformed the constructs into *B. napus* DH12075 using the C58C1 strain. In four tested independent hairy root lines, application of 1 µM NAA or 1 µM IAA elicited a measurable auxin signaling response after 6 and 24 h, as evidenced by significant increase in fluorescence. The response exhibited a broad spatial distribution and was more pronounced in the VENUS fusions (Supplementary Fig. S11 – S12).

Auxin plays a key role in integrating abiotic stress signals and mediating plant responses (Jing et al. 2023). To investigate auxin output in *B. napus* under stress conditions, hairy roots expressing DR5cc-Ω-VENUS and DR5cc-Ω-mCherry were treated with 150 mM mannitol or 75 mM NaCl to mimic drought and salinity, respectively. Fluorescence intensity was quantified at 1, 6, 12, and 24 h to monitor temporal dynamics. Both treatments induced a transient auxin response in the root apex (Fig. 4, Supplementary Fig. S13). No significant changes relative to mock-treated controls were observed after 1 h of application of the stress inducers, whereas fluorescence intensity increased significantly at 6 and 12 h and returned to control levels after 24 h of treament. Such temporal dynamic pattern was reproducible across independent transgenic lines and reporter variants (Fig. 4, Supplementary Fig. S13 – S15).

**Fig. 4.**
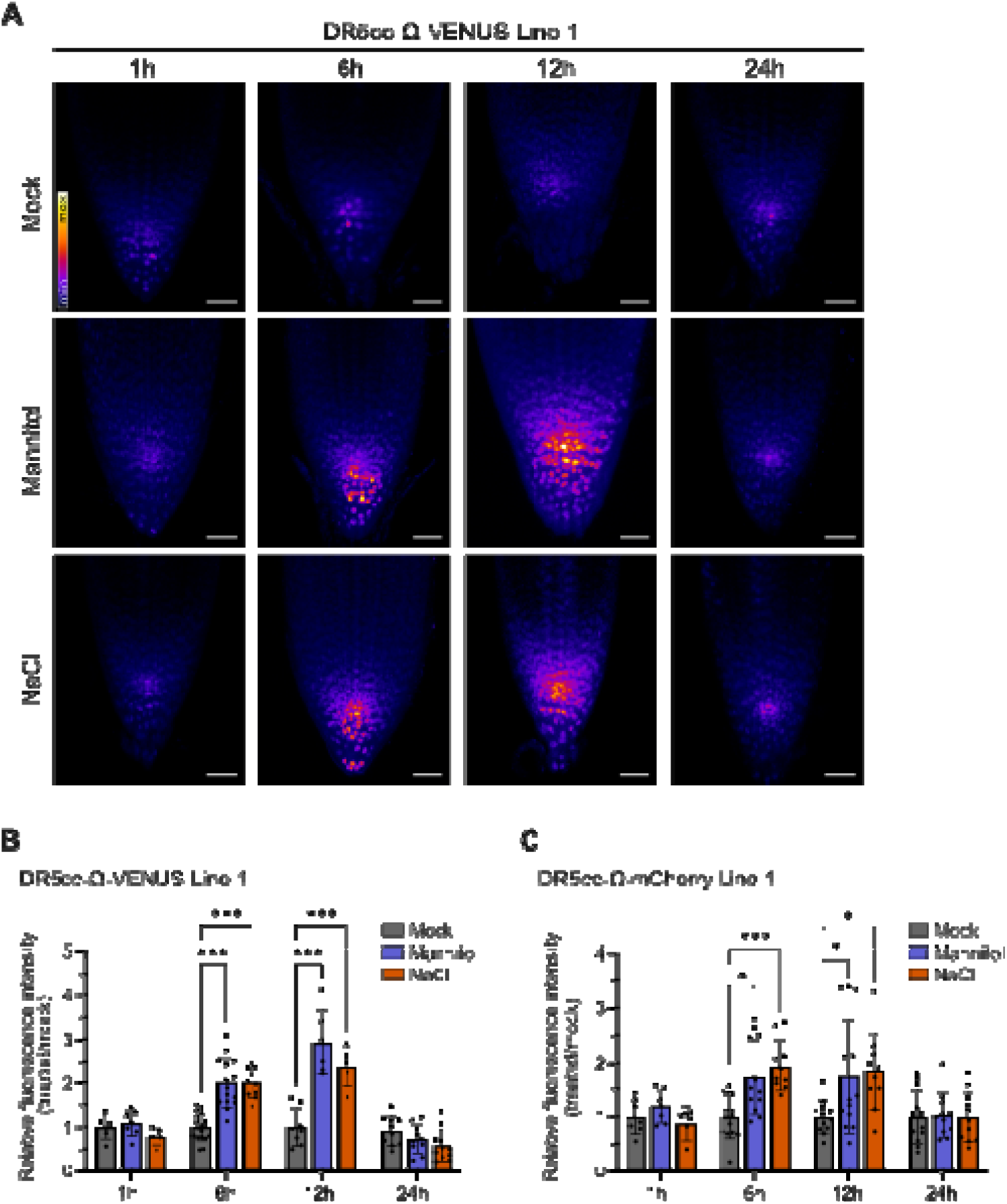
Stress-induced auxin signaling dynamics in *B. napus* hairy roots. **(A)** Maximum intensity projections of root tips expressing DR5cc-Ω-VENUS under mock and abiotic stress conditions (150 mM mannitol or 75 mM NaCl), imaged at 1, 6, 12, and 24 h of treatment. Scale bars represent 40 µm. **(B, C)** Quantification of relative VENUS **(B)** and mCherry **(C)** fluorescence intensities in mock- and stress-treated hairy roots at indicated time points. Error bars indicate SD (n = 5-19 roots). For each time point, values are normalized to mock. Statistical comparisons (one-way ANOVA with Dunnett’s post-hoc test) were performed independently within each time point to compare stress treatments against the corresponding mock; (*P < 0.05, ***P < 0.001); unannotated comparisons within a time point indicate no statistically significant difference (P ≥ 0.05). Images corresponding to (C) are presented in Supplementary Fig. S13

### Concurrent analysis of cytokinin and auxin signaling in tissue culture

We previously demonstrated the responsiveness of the *Arabidopsis thaliana*-derived TCSv2 reporter (Steiner et al. 2020) to application of cytokinin and abiotic stress in hairy roots of *B. napus* cv. Darmor and *B. rapa* R-o-18 (Nicolas Mala et al. 2024). In this study, we introduced the same construct, TCSv2-VENUS, into *B. napus* DH12075 and regenerated plants from the transformed hairy roots. The cytokinin TCSv2-VENUS reporter was functional in the roots of regenerated plants, as evidenced by its induction following a 2 h treatment with 5 µM BAP (Fig. 5A, B).

**Fig. 5.**
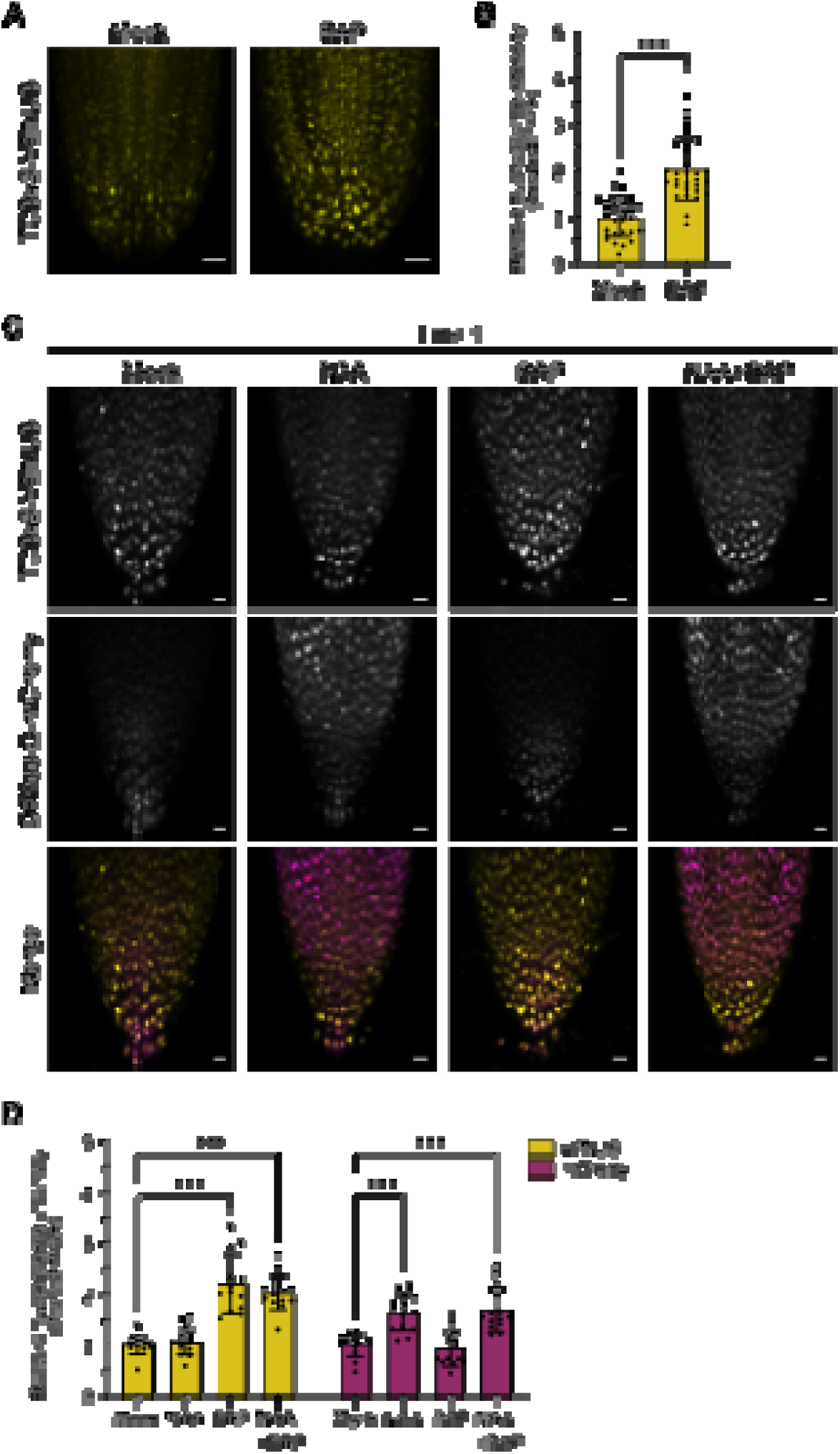
Cytokinin and auxin reporter activity in *B. napus*. (A) Maximum intensity projections of mock- and BAP-treated (5 µM, 2 h) roots from regenerated plants expressing TCSv2-VENUS. Scale bars represent 50 µm. (B) Quantification of relative VENUS fluorescence intensity in mock- and BAP-treated root tips. Error bars indicate SD (n = 12-30 roots); ***P < 0.001; Welch’s t-test. Values are normalized to mock. (C) Maximum intensity projections of hairy roots co-expressing TCSv2-VENUS and DR5cc-Ω-mCherry treated with mock, NAA (1 µM, 24 h), and / or BAP (5 µM, 2 h). Images are shown as VENUS (top row), mCherry (middle row) and merged channels (bottom row: magenta color for DR5cc-Ω-mCherry and yellow for TCSv2-VENUS). Scale bars represent 20 µm. (D) Quantification of relative VENUS (left) and mCherry (right) fluorescence intensities in mock- and hormone-treated hairy roots. Error bars indicate SD (n = 7-14 roots). For each channel, values are normalized to mock. Statistical comparisons (one-way ANOVA with Dunnett’s post-hoc test) were performed independently within each channel to compare hormonal treatments against the corresponding mock; (***P < 0.001); unannotated comparisons within a time point indicate no statistically significant difference (P ≥ 0.05)

The TCSv2-VENUS plants were subsequently transformed using the *Agrobacterium* strain C58C1 carrying the DR5cc-Ω-mCherry auxin reporter construct. Two independent hairy root lines were treated with NAA (1 µM, 24 h) and / or BAP (5 µM, 2 h), and mCherry and VENUS fluorescence intensities were quantified in root tips (Fig. 5C, D, Supplementary Fig. S16). In both lines, only the reporter corresponding to the applied hormone (VENUS for BAP and mCherry for NAA) showed significant increase in fluorescence and broader signal distribution, demonstrating that both hormonal responses can be simultaneously monitored in *B. napus* hairy roots.

### DII-based reporters for oilseed rape

We developed a degron-based quantitative DII (qDII) reporter system for oilseed rape. The DII domains of two *B. napus* Aux/IAA proteins (IAA2 and IAA17) were synthesized in frame with the VENUS fluorescent protein. Each construct also includes TagBFP, a stable blue fluorescent protein that is not degraded and serves as a reference for signal normalization. The two fluorescent proteins are separated by a 2A peptide, and the expression of the constructs is driven by the full CaMV 35S promoter. The DII domains contain not only the degron motif (VGWPPVRS[Y/S]R) but also the upstream region with the KR motif and the downstream degron tail (Fig. 6A), as these elements have been shown to be important for auxin-dependent degradation (Figueiredo and Strader 2022). Considering the A and C subgenomes and multiple gene homologs, the *B. napus* sequences encoding the DII domains were selected based on the presence of characteristic motifs (Supplementary Fig. S3) and supporting expression data (Chao et al. 2020).

**Fig. 6.**
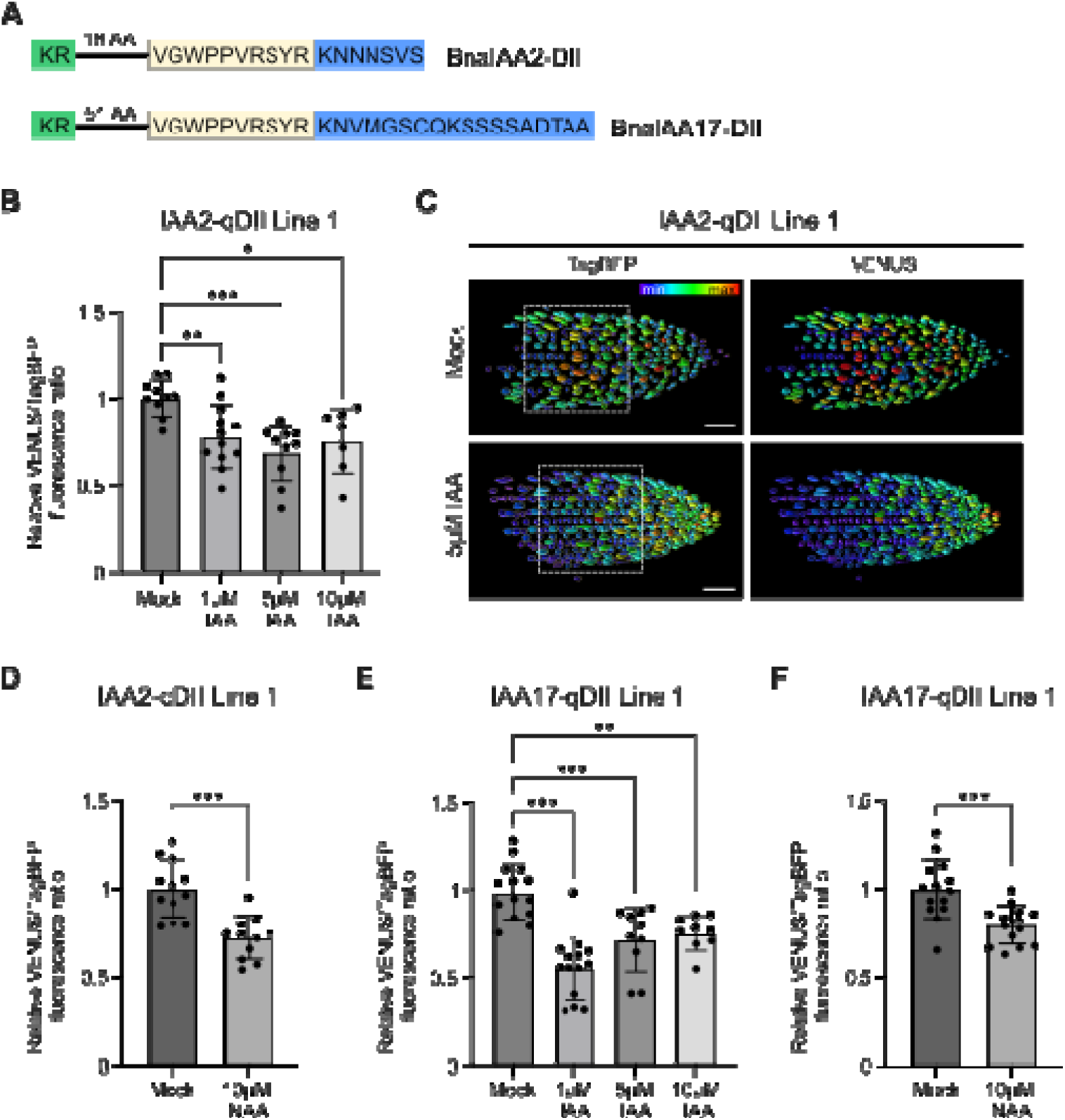
qDII reporters are auxin responsive in *B. napus* hairy roots. (A) Schematic representation of the selected DII domains. (B–D) Response of IAA2-qDII Line 1 to auxin treatment. (B) Quantification of the nuclear VENUS/TagBFP fluorescence intensity ratio following treatment with different IAA concentrations (2 h). (C) Maximum intensity projections of hairy root tips under mock and IAA (5 µM, 2 h) conditions, showing measured signal intensities of TagBFP (left) and VENUS (right). The boxed region indicates the area used for nuclear signal quantification. Images were generated using IMARIS software. Scale bars represent 50 µm. (D) Quantification of the nuclear VENUS/TagBFP fluorescence intensity ratio following NAA treatment (10 µM, 2 h). (E–F) Response of IAA17-qDII Line 1 to auxin treatment. Quantification of the nuclear VENUS/TagBFP fluorescence intensity ratio following treatment with different IAA concentrations (2 h) (E) or NAA (10 µM, 2 h) (F). Statistical significance was assessed by one-way ANOVA followed by Dunnett’s post-hoc test for IAA treatment (*P < 0.05, **P < 0.01, ***P < 0.001) and Welch’s t-test for NAA treatment (***P < 0.001). Values are normalized to the corresponding mock controls. Error bars represent SD (n = 7 – 14 roots).

Hairy roots carrying the IAA2-qDII or IAA17-qDII constructs were treated with IAA (1 µM, 5 µM, or 10 µM for dose-response experiments) or NAA (10 µM) for 2 hours, followed by imaging of epidermal cells within the root transition zone. All treatments resulted in a significant decrease in the nuclear VENUS/TagBFP ratio across all tested lines (Line 1 of each construct is shown in Fig. 6; Line 2 in Supplementary Fig. S17 – S19). In contrast, nuclear TagBFP fluorescence remained stable upon NAA treatment, confirming the stability of the reference channel (Supplementary Fig. S20). These results demonstrate that the qDII system is functional and enables quantitative readout of auxin responses in *B. napus* hairy roots.

## Discussion

*Agrobacterium* strains carrying hairy root–inducing (Ri) plasmids trigger abnormal root formation in host plants by transferring T-DNA into the plant genome at wound sites. Expression of T-DNA genes, particularly the *rol* loci, drives hairy root development (Christey 2001). Agrobacterial strains harboring both an Ri plasmid and an engineered binary vector are widely employed for the delivery of foreign DNA into plant cells. Hairy root tissue culture expressing DR5 transgene has previously been demonstrated to be a suitable system for monitoring auxin signaling across various species (Demina et al. 2019; Shrestha et al. 2021; Flores-Moreno et al. 2026).

Earlier studies investigating auxin signaling in *B. napus* employed DR5 reporters containing the TGTCGG and TGTCTC AuxRE variants during plant development and the early stages of microspore embryogenesis (Dubas et al. 2014; Zhang et al. 2025). Because different TGTCNN AuxRE variants have been shown to exhibit dose-dependent responses to auxin while differing markedly in their response strength in *Arabidopsis* (Lieberman-Lazarovich et al. 2019), we designed a reporter containing the TGTCCC AuxRE motif, which has been associated predominantly with auxin-responsive genes based on gene ontology analysis. In addition, we introduced variable spacer sequences between AuxRE motifs to avoid a highly repetitive synthetic promoter structure. Both the DR5cc reporter and the *B. napus*-derived BIP3 reporter, which is based on promoter fragments of auxin-responsive genes, demonstrated functionality and responsiveness to auxin in both hairy root cultures and regenerated plants. When used to drive GUS expression, these reporters, like conventional GUS-based reporters, enable spatial analysis of auxin responses within plant tissues during development. In hairy root cultures generated by two different *Agrobacterium* strains (the cucumopine-type strain K599 and the transconjugant C58C1 carrying the agropine-type *pRiA4*), auxin signaling remained comparable. Differences were observed only in the growth of the hairy root lines, with those derived from K599 exhibiting slower growth than those obtained after transformation with C58C1. This observation is consistent with a previous study indicating that T-DNA-encoded genes acquired from the K599 strain negatively affect root growth under both normal and stress conditions in tobacco regenerants (Shvets et al. 2024).

In regenerated plants, attempts to segregate the Ri plasmid T-DNA from the T-DNA of the binary vector were unsuccessful. A similar limitation was observed in additional reporter lines developed in our laboratory (data not shown). One possible explanation is the use of the transconjugant C58C1 strain, which may exhibit an increased propensity for co-integration of multiple T-DNAs (specifically those derived from the *Ri* plasmid and the binary vector) into the same genomic locus. Previous studies using *A. tumefaciens* have investigated co-transformation frequencies of two T-DNAs. With the C58C1 strain, the described co-transformation system favored genetically linked integration of the two T-DNAs (De Block and Debrouwer 1991), whereas a study employing the LBA4404 strain did not observe preferential integration into linked loci (Komari et al. 1996). As our optimized protocol utilized the transconjugant C58C1 strain carrying pRiA4, this phenomenon may be associated with the *Agrobacterium* strain used. Alternatively, T-DNA integration patterns are known to be influenced by other factors, including plant species, the cell type targeted for transformation, vector design, and the transformation method employed (De Buck et al. 2009; Al Nuaimi et al. 2025). In our case, the injection-based transformation method may also contribute to this effect.

Despite the confirmed presence of Ri plasmid T-DNA in regenerated oilseed rape plants exhibiting the Ri phenotype, auxin signaling in these lines was comparable to that observed in Arabidopsis plants lacking *Ri* T-DNA (Aloni et al. 2006; Smolko et al. 2021). The Ri phenotype in oilseed rape primarily affects plant stature and leaf morphology, as previously documented for *B. napus* DH12075 (Jedličková et al. 2022b, 2023). Therefore, hairy-root-derived auxin reporter regenerants may not be suitable for experiments addressing these traits. In seedlings, however, the Ri phenotype is not readily distinguishable from the wild type, and flowers and embryos also appear comparable to those of wild-type plants (Figure 3). Thus, although genes located on the Ri plasmid T-DNA can modulate auxin homeostasis and enhance plant sensitivity to this hormone (Maurel et al. 1991; Mauro et al. 2017), oilseed rape regenerants remain suitable for auxin signaling studies.

Fluorescent reporters can be used to monitor dynamic changes in hormone signaling. They are suitable for studying responses to both endogenous signaling events and the application of exogenous hormones. Previously, we demonstrated stress-induced changes in cytokinin signaling using a TCS-based reporter in Brassica hairy roots (Nicolas Mala et al. 2024). Here, we extend this approach to auxin by analyzing *B. napus* hairy roots expressing DR5 fluorescent reporters. We observed a transient increase in DR5 signal in root tips exposed to osmotic and salt stress, suggesting a temporary change in auxin responsiveness. In Arabidopsis, osmotic and salt stress alter local auxin maxima in roots within hours (Smolko et al. 2021). Similarly, salinity induces rapid redistribution of auxin signaling output in root tips during halotropic growth (Galvan-Ampudia et al. 2013). These observations are consistent with the dynamic nature of auxin distribution in plant tissues and its role in adaptive growth responses (Jing et al. 2023; Vanneste et al. 2025). Our results further support the use of fluorescent DR5-based reporters and demonstrate their applicability for monitoring early stress responses in crop species.

DR5 reporters reflect downstream signaling output rather than auxin levels themselves. To more directly assess cellular auxin distribution, reporters based on the auxin-sensitive domain (DII) have been developed as auxin input reporters (Jedličková et al. 2022a). Previous studies have shown that regions flanking the degron modulate degradation dynamics (Moss et al. 2015; Figueiredo and Strader 2022). Therefore, we developed a qDII reporter incorporating the degron together with key flanking regions, including the degron tail and proximal KR motif, derived from two Aux/IAA proteins, IAA2 and IAA17. Using a hairy root transformation system, we demonstrated the functionality of the qDII reporter in oilseed rape.

In conclusion, we developed auxin input and output reporters for oilseed rape, enabling visualization of auxin signaling in both hairy root systems and whole plants. These reporters provide a versatile platform for the rapid evaluation of auxin dynamics in transformed roots and for monitoring spatial auxin patterns *in planta*, expanding the toolkit for studying hormone response in crop species.

## Supplementary material

**Supplementary Fig. S1** Schematic representation of the constructs prepared for this study. Addgene plasmids used for MoClo cloning are indicated

**Supplementary Fig. S2** Promoter fragments of *BnaIAA* genes used in the BIP3 reporter construct. AuxRE motifs in the forward orientation (TGTCCC) are indicated by green boxes, whereas AuxRE motifs in the reverse orientation (GGGACA) are indicated by blue boxes

**Supplementary Fig. S3** *BnaIAA* genes selected for qDII reporter construction. The coding regions of *BnaA01g24190D* (A; *IAA2*) and *BnaA08g27770D* (B; *IAA17*), together with their corresponding amino acid sequences, are shown. KR motif is highlighted in green, the degron motif in yellow, and the degron tail in blue

**Supplementary Fig. S4** Sequence alignment of the two *ACC* gene copies in *B. napus* DH12075. Amplification primers for the 0.6 kb fragment are highlighted in green, whereas primers and probes specific for the *ACC*-A copy used in qPCR are highlighted in blue and yellow, respectively

**Supplementary Fig. S5** GUS expression in hairy roots carrying reporter constructs driven by different promoters. Hairy roots expressing (**A**) the minimal CaMV 35S promoter-GUS reporter, (**B**) the full CaMV 35S promoter-GUS reporter, (**C**) the DR5cc-GUS reporter, Line 2 (three transgene copies), (**D**) BIP3-GUS, Line 2 (five transgene copies), and (**E**) BIP3-Ω-GUS, Line 2 (two transgene copies). Hairy roots were either mock-treated or treated with 0.1 μM or 1 μM NAA for 12 h prior to GUS staining. Scale bars represent 500 µm

**Supplementary Fig. S6** Growth of hairy roots induced by the C58C1 or K599 *Agrobacterium* strains. (A) Relative root length (% of the initial root length at day 0) was measured after 8 and 16 days of culture in four lines transformed by C58C1 and four lines transformed by K599. Three independent biological replicates were performed, and the data are presented as mean values with standard deviation. (B) Representative images of hairy root lines at the initial time point and after 8 days of culture. Scale bar represents 1 mm

**Supplementary Fig. S7** Histochemical GUS staining of different tissues of *B. napus* transgenic plants. Independent transgenic lines from (A) DR5cc-GUS (+/- Ω) and (B) BIP3-GUS (+/- Ω) are presented. Scale bars represent 2 mm (seedlings, flowers), 200 µm (root tips), and 500 µm (embryos)

**Supplementary Fig. S8** Histochemical GUS staining of *B. napus* flowers expressing auxin reporters after NAA treatment. Flowers were sprayed with water (Mock) or 50 µM NAA in water, and stained after 24 h. Scale bars represent 2 mm

**Supplementary Fig. S9** Histochemical GUS staining of roots from regenerated transgenic *B. napus* plants following NAA treatment. Representative images showing GUS activity in root tips expressing DR5cc-GUS (+/- Ω) and BIP3-GUS (+/- Ω) under mock control conditions or following treatment with 0.1 and 1 µM NAA for 3 h. Scale bars represent 200 µm

**Supplementary Fig. S10** Histochemical GUS staining of roots from regenerated transgenic *B. napus* plants following IAA treatment. Representative images showing GUS activity in root tips expressing DR5cc-GUS (+/- Ω) and BIP3-GUS (+/- Ω) under mock control conditions or following treatment with 0.1 and 1 µM IAA for 3 h. Scale bars represent 200 µm

**Supplementary Fig. S11** The DR5cc reporter is responsive to NAA treatment in *B. napus* hairy roots. (A) Maximum intensity projections of root tips from independent lines expressing DR5cc-Ω-VENUS or DR5cc-Ω-mCherry treated with mock or 1 µM NAA, and imaged after 6 or 24 h. Scale bars represent 40 µm. (B) Quantification of relative VENUS and mCherry fluorescence intensities in mock- and NAA-treated hairy roots at indicated time points. Error bars indicate SD (n = 7-14 roots). For each time point, values are normalized to the corresponding mock. Statistical comparisons (one-way ANOVA with Dunnett’s post-hoc test) were performed independently within each time point to compare stress treatments against the corresponding mock (**P < 0.01, ***P < 0.001)

**Supplementary Fig. S12** The DR5cc reporter is responsive to IAA treatment in *B. napus* hairy roots. (A) Maximum intensity projections of root tips from independent lines expressing DR5cc-Ω-VENUS or DR5cc-Ω-mCherry treated with mock or 1 µM IAA, and imaged after 6 or 24 h. Scale bars represent 40 µm. (B) Quantification of relative VENUS and mCherry fluorescence intensities in mock- and IAA-treated hairy roots at indicated time points. Error bars indicate SD (n = 6-10 roots). For each time point, values are normalized to the corresponding mock. Statistical comparisons (one-way ANOVA with Dunnett’s post-hoc test) were performed independently within each time point to compare stress treatments against the corresponding mock (*P < 0.5, **P < 0.01, ***P < 0.001)

**Supplementary Fig. S13** Stress-induced auxin signaling dynamics in hairy roots expressing DR5cc-Ω-mCherry (Line 1). Maximum intensity projections of root tips treated with mock and abiotic stress (150 mM mannitol or 75 mM NaCl), and imaged at 1, 6, 12, and 24 h. Scale bars represent 40 µm. Quantification is shown in Fig. 4.

**Supplementary Fig. S14** Stress-induced auxin signaling dynamics in hairy roots expressing DR5cc-Ω-VENUS (Line 2). (A) Maximum intensity projections of root tips treated with mock and abiotic stress (150 mM mannitol or 75 mM NaCl), and imaged at 1, 6, 12, and 24 h. Scales bars represent 40 µm. (B) Quantification of relative VENUS fluorescence intensity in mock- and stress-treated hairy roots at indicated time points. Error bars indicate SD (n = 6-18 roots). For each time point, values are normalized to the corresponding mock. Statistical comparisons (one-way ANOVA with Dunnett’s post-hoc test) were performed independently within each time point to compare stress treatments against the corresponding mock (**P < 0.01, ***P < 0.001); unannotated comparisons within a time point indicate no statistically significant difference (P ≥ 0.05)

**Supplementary Fig. S15** Stress-induced auxin signaling dynamics in hairy roots expressing DR5cc-Ω-mCherry (Line 2). (A) Maximum intensity projections of root tips treated with mock and abiotic stress (150 mM mannitol or 75 mM NaCl), and imaged at 1, 6, 12, and 24 h. Scale bars represent 40 µm. (B) Quantification of relative mCherry fluorescence intensity in mock- and stress-treated hairy roots at indicated time points. Error bars indicate SD (n = 7-16 roots). For each time point, values are normalized to the corresponding mock. Statistical comparisons (one-way ANOVA with Dunnett’s post-hoc test) were performed independently within each time point to compare stress treatments against the corresponding mock (*P < 0.05; **P < 0.01, ***P < 0.001); unannotated comparisons within a time point indicate no statistically significant difference (P ≥ 0.05)

**Supplementary Fig. S16** Cytokinin and auxin reporter activity in *B. napus* hairy roots, transgenic Line 2. (A) Maximum intensity projections of hairy roots co-expressing TCSv2-VENUS and DR5-mCherry treated with mock, NAA (1 µM, 24 h), and/or BAP (5 µM, 2 h). Images are shown as VENUS (top row), mCherry (middle row) and merged channels (bottom row: magenta color for DR5cc-Ω-mCherry and yellow for TCSv2-VENUS). Scale bars represent 20 µm. (B) Quantification of relative VENUS (left) and mCherry (right) fluorescence intensities in mock- and hormone-treated hairy roots. Error bars indicate SD (n = 7-9 roots). For each channel, values are normalized to the corresponding mock. Statistical comparisons (one-way ANOVA with Dunnett’s post-hoc test) were performed independently within each channel to compare hormonal treatments against the corresponding mock (***P < 0.001); unannotated comparisons within a time point indicate no statistically significant difference (P ≥ 0.05)

**Supplementary Fig. S17.** Auxin responsiveness of the IAA2-qDII reporter in *B. napus* hairy roots following NAA treatment. (A) Representative images of epidermal cells from IAA2-qDII Line 2 under mock and NAA (10 µM, 2 h) conditions, showing TagBFP (left) and VENUS (right) fluorescence. (B) Quantification of the nuclear VENUS/TagBFP fluorescence intensity ratio for the conditions shown in (A) (**P < 0.01, Welch’s t-test). Values are normalized to the corresponding mock. Error bars indicate standard deviation (SD; n = 13 roots per treatment). (C) Representative images of epidermal cells from IAA2-qDII Line 1 under mock and NAA (10 µM, 2 h) conditions, showing TagBFP (left) and VENUS (right) fluorescence. The corresponding quantification of the nuclear VENUS/TagBFP fluorescence intensity ratio is presented in Figure 6D. Scale bars represent 20 μm.

**Supplementary Fig. S18.** Auxin responsiveness of the IAA17-qDII reporter in *B. napus* hairy roots following NAA treatment. (A) Representative images of epidermal cells from IAA17-qDII Line 2 under mock and NAA (10 µM, 2 h) conditions, showing TagBFP (left) and VENUS (right) fluorescence. (B) Quantification of the nuclear VENUS/TagBFP fluorescence intensity ratio for the conditions shown in (A) (***P < 0.001, Welch’s t-test). Values are normalized to the corresponding mock. Error bars indicate standard deviation (SD; n = 18 − 21 roots). (C) Representative images of epidermal cells from IAA17-qDII Line 1 under mock and NAA (10 µM, 2 h) conditions, showing TagBFP (left) and VENUS (right) fluorescence. The corresponding quantification of the nuclear VENUS/TagBFP fluorescence intensity ratio is presented in Figure 6F. Scale bars represent 20 μm.

**Supplementary Figure S19**. IAA responsiveness of IAA2-qDII Line 2 and IAA17-qDII Line 2 in B. napus hairy roots. Maximum intensity projections of hairy root tips from (A) IAA2-qDII Line 2 and (B) IAA17-qDII Line 2 under mock and IAA (5 µM, 2 h) treatment conditions, showing measured signal intensities of TagBFP (left) and VENUS (right). The boxed region indicates the area used for nuclear signal quantification. Images were generated using IMARIS software. Scale bars represent 50 µm. Quantification of the nuclear VENUS/TagBFP fluorescence intensity ratio for (C) IAA2-qDII Line 2 and (D) IAA17-qDII Line 2. Statistical significance was assessed by one-way ANOVA followed by Dunnett’s post hoc test (*P < 0.05, **P < 0.01, ***P < 0.001). Values are normalized to the corresponding mock controls. Error bars indicate standard deviation (SD; n = 9 – 17 roots).

**Supplementary Fig. S20** Stability of the TagBFP fluorescence in qDII constructs. Quantification of relative TagBFP fluorescence intensity in independent hairy root lines expressing IAA2 / IAA17-qDII treated with mock or NAA (10 µM, 2 h). Error bars indicate SD (n = 11-21 roots). No significant differences were observed in treated samples compared with the mock control (Welch’s t-test)

**Supplementary Table S1.** List of oligonucleotides.

## Statements and Declarations

### Author Contribution Statement

VJ and HSR conceived and designed the research. VJ, VP, MŠt, MZ, and MSe conducted experiments. VJ, VP, and MZ analyzed data. VJ, VP and HSR wrote the manuscript. All authors read and approved the manuscript.

### Data availability

All data supporting the findings of this study are available in the article, in its supplementary information files, and in Zenodo repository (https://doi.org/10.5281/zenodo.21030825).

## Acknowledgements and Funding

We acknowledge the Core Facility CELLIM supported by the Czech-BioImaging large RI project (LM2023050 funded by MEYS CR) and the Core Facility Plant Sciences of CEITEC Masaryk University for their support in obtaining scientific data presented in this paper. This work was funded by the Czech Science Foundation project GA23-06140S and supported by the project TowArds Next GENeration Crops (TANGENC), reg. no. CZ.02.01.01/00/22_008/0004581 of the ERDF Programme Johannes Amos Comenius.

