## Supplemental Figures for "Development of auxin reporters in oilseed rape (*Brassica napus*)"

Plant Cell Reports

**Supplementary Figure S1.** Schematic representation of the constructs prepared for this study.

**Supplementary Figure S2.** Promoter fragments of *BnaIAA* genes used in the BIP3 reporter construct.

**Supplementary Figure S3.** *BnaIAA* genes selected for qDII reporter construction.

**Supplementary Figure S4.** Sequence alignment of the two *ACC* gene copies in *B. napus* DH12075.

**Supplementary Figure S5.** GUS expression in hairy roots carrying reporter constructs driven by different promoters.

**Supplementary Figure S6.** Growth of hairy roots induced by the C58C1 or K599 *Agrobacterium* strains.

**Supplementary Figure S7.** Histochemical GUS staining of different tissues of *B. napus* transgenic plants.

**Supplementary Figure S8.** Histochemical GUS staining of *B. napus* flowers expressing auxin reporters after NAA treatment.

**Supplementary Figure S9.** Histochemical GUS staining of roots from regenerated transgenic *B. napus* plants following NAA treatment.

**Supplementary Figure S10.** Histochemical GUS staining of roots from regenerated transgenic *B. napus* plants following IAA treatment.

**Supplementary Figure S11.** The DR5cc reporter is responsive to NAA treatment in *B. napus* hairy roots.

**Supplementary Figure S12.** The DR5cc reporter is responsive to IAA treatment in *B. napus* hairy roots.

**Supplementary Figure S13.** Stress-induced auxin signaling dynamics in hairy roots expressing DR5cc-Ω-mCherry (Line 1).

**Supplementary Figure S14.** Stress-induced auxin signaling dynamics in hairy roots expressing DR5cc-Ω-VENUS (Line 2).

**Supplementary Figure S15.** Stress-induced auxin signaling dynamics in hairy roots expressing DR5cc-Ω-mCherry (Line 2).

**Supplementary Figure S16.** Cytokinin and auxin reporter activity in *B. napus* hairy roots, transgenic Line 2.

**Supplementary Figure S17.** Auxin responsiveness of the IAA2-qDII reporter in *B. napus* hairy roots following NAA treatment.

**Supplementary Figure S18.** Auxin responsiveness of the IAA17-qDII reporter in *B. napus* hairy roots following NAA treatment.

**Supplementary Figure S19.** IAA responsiveness of IAA2-qDII Line 2 and IAA17-qDII Line 2 in *B. napus* hairy roots.

**Supplementary Figure S20.** Stability of the TagBFP fluorescence in qDII constructs.

**Supplementary Table S1.** List of oligonucleotides.

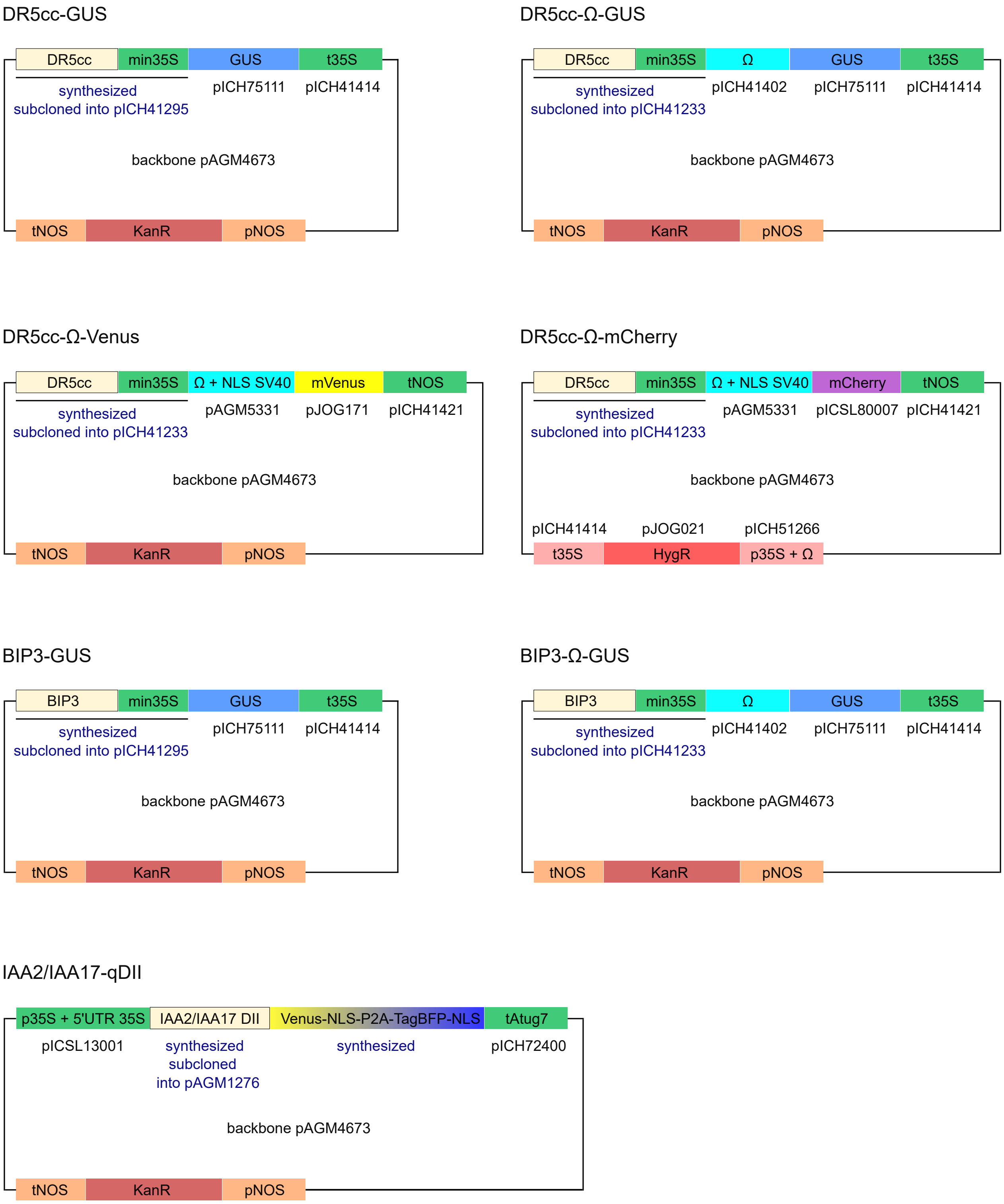


**Supplementary Figure S1**. Schematic representation of the constructs prepared for this study. Addgene plasmids used for MoClo cloning are indicated.

```
                    pBnaIAA1
BnaC08g09640D    AAAGTAACGCGTCCAAATATCTCAGTGTCCCATCTTTGTCCCCTTGCCTCTC
BnaA08g30190D    AAAGTAACGCGTCCAAATATCTCAGTGTCCCATCTTTGTCCCCTTGCCTCTC
                 ****************************************************

                    pBnaIAA2
BnaA05g16700D    ACGAGTCCACATGGGCGGCCATAGCGTTTGTGTCCTACCTTTGTCCCCTTGC
BnaC05g29330D    TCGAGTCCACATGGGCGGCCATAGCGTTTGTGTCCTACCTTTGTCCCCTTGC
BnaA01g24190D    TCGAGTCCACATGGGCGGCCATAGCGTTTGTGTCCCACCTTTGTCCCCTTGC
BnaC01g31190D    TCGAGTCCACATGGGCGGCCATAGCGTTTGTGTCCCACCTTTGTCCCCTTGC
BnaA03g36940D    TCGAGTTCACATGGACGGCCAAAGCGTTTATGTCCCACCTTTGTCCCCTTGC
BnaC03g43120D    TCGAGTTCACATGGACGGCCAAAGCGTTTATGTCCCACCTTTGTCCCCTTGC
                  ***** ******* ****** ******* ***** ****************

                    pBnaIAA4
BnaA09g16590D    TTTGGCTATAGATGAAAGTGTCCCACGAATGTCCCCAAAATGATGGGACAC
BnaCnng63870D    TTTGGCTATAGATGAAAGTGTCCCACGAATGTCCCCAAAATGATGGGACAC
                 ***************************************************
```

**Supplementary Figure S2**. Promoter fragments of *BnaIAA* genes used in the BIP3 reporter construct. AuxRE motifs in the forward orientation (TGTCCC) are indicated by green boxes, whereas AuxRE motifs in the reverse orientation (GGGACA) are indicated by blue boxes.

## A

```
atggcttacgagaaagtcaatgagcttaaccttaaggacacagagcttcgtcttggatta
 M  A  Y  E  K  V  N  E  L  N  L  K  D  T  E  L  R  L  G  L
cccggaacggagcaagataaggaagaacaagaggtttcttgcgttagaagcaacaagcgt
 P  G  T  E  Q  D  K  E  E  Q  E  V  S  C  V  R  S  N  K  R
caactacagagcgataacgaggaagaatctacacttcctacgaaaactcaaatcgttggt
 Q  L  Q  S  D  N  E  E  E  S  T  L  P  T  K  T  Q  I  V  G
tggcctccggtgagatcttaccgtaaaaacaacaacagtgtgagctatgtgaaagtgagt
 W  P  P  V  R  S  Y  R  K  N  N  N  S  V  S  Y  V  K  V  S
atggatggagctccataccttaggaaaatagatctcaagacatacaaaaactatccagag
 M  D  G  A  P  Y  L  R  K  I  D  L  K  T  Y  K  N  Y  P  E
cttctcaaggcattagagaacctgttcaagttcacgatcggtgaatacaacgaaagagaa
 L  L  K  A  L  E  N  L  F  K  F  T  I  G  E  Y  N  E  R  E
ggatacaaaggatctggagttgtaccaacgtacgaggataaagatggagattggatgttg
 G  Y  K  G  S  G  V  V  P  T  Y  E  D  K  D  G  D  W  M  L
gttggtgatgttccatgggatatgttctcttcctcttgtaagagactcaggatcatgaaa
 V  G  D  V  P  W  D  M  F  S  S  S  C  K  R  L  R  I  M  K
ggatctgatgctcttgctttggactcggccttatga
 G  S  D  A  L  A  L  D  S  A  L  *
```

## B

```
atgatgggcagtgttgggctgaatctgagggagactgagctgtgtcttggtcttcccggt
 M  M  G  S  V  G  L  N  L  R  E  T  E  L  C  L  G  L  P  G
ggtgatacggcgtccccgttgactggaaccaagagaggattctcagagacggttgatctg
 G  D  T  A  S  P  L  T  G  T  K  R  G  F  S  E  T  V  D  L
aagctgaatctgaacaatgaacctgaaagcaaggaaggatctaagagccacgacgtcgtg
 K  L  N  L  N  N  E  P  E  S  K  E  G  S  K  S  H  D  V  V
agtgctatttccaaggaaaagagttcatgtaccaaagatccaaccaagcctcctgccaag
 S  A  I  S  K  E  K  S  S  C  T  K  D  P  T  K  P  P  A  K
gcacaagttgtgggatggccaccggtgagatcataccggaagaacgtgatgggttcatgc
 A  Q  V  V  G  W  P  P  V  R  S  Y  R  K  N  V  M  G  S  C
caaaaatcaagcagtagcgcggacacggcggcgtttgtgaaggtgtcgatggacggagca
 Q  K  S  S  S  S  A  D  T  A  A  F  V  K  V  S  M  D  G  A
ccgtacttgaggaaaatcgacttgaagatgtataagagctacgacgaactctctgacgct
 P  Y  L  R  K  I  D  L  K  M  Y  K  S  Y  D  E  L  S  D  A
ttgtccaacatgttcagctcttttaccatgggaaaaaatggaggagaagaaggaatgata
 L  S  N  M  F  S  S  F  T  M  G  K  N  G  G  E  E  G  M  I
gacttcatgaatgagaggaaagtaatggatacagtgagtagttgggactatgttccctct
 D  F  M  N  E  R  K  V  M  D  T  V  S  S  W  D  Y  V  P  S
tatgaagacaaagacggtgattggatgctcgtcggcgacgtcccttggccaatgttcgtt
 Y  E  D  K  D  G  D  W  M  L  V  G  D  V  P  W  P  M  F  V
gatacatgcaagcgtttacgtctcatgaagggatctgacgccattggtctcgctcctaga
 D  T  C  K  R  L  R  L  M  K  G  S  D  A  I  G  L  A  P  R
gcaatggagaagtgcaagagtagagcttga
 A  M  E  K  C  K  S  R  A  *
```

**Supplementary Figure S3**. *BnaIAA* genes selected for qDII reporter construction. The coding regions of *BnaA01g24190D* (**A**; IAA2) and *BnaA08g27770D* (**B**; IAA17), together with their corresponding amino acid sequences, are shown. KR motif is highlighted in green, the degron motif in yellow, and the degron tail in blue.

```
                      ACC_amp_F
Bna_acc-C       ATCGCTGGAGTTTCTGCCGAATATAATAAGTGCAGCA---CTGACAGAAACAAACCACAG
Bna_acc-A       ATCGCTGGAGTTTCTGCCGAATATAATAACTGCAGCAGCACTGACAGAAACAAACCACAT
                ***************************** *******   *******************

Bna_acc-C       CGACTATGA------------ATCTCCTTTATCTGGCAATATAATGCACATTGCTATTGT
Bna_acc-A       CGACTATGGTGAATCTGCCGGATCTCCTCTATGTGGCAATATAATGCACATTGCTGTTTT
                ********             ******* *** ********************** ** *

Bna_acc-C       GGGCATGAGTCTTCTTCAGGACAGGTACTTGACACAAGCCTGTGACGTTTTCAAACTAAG
Bna_acc-A       GG------------------ACAGGTACTTCACACAGCC---TGACGGTTTCAAACTAGC
                **                  ********** *****  *   ***** **********

Bna_acc-C       GTAGGTGCTAGA-TATATCGTCTAACCATATCTGTTAACGCTGCTGCTTTAATTTGTTCG
Bna_acc-A       TTAGGTGCTAAGCTATATTGTCTAACCTTATCTATTAACACTGCTGCTTTAATATGTTCG
                 *********   ***** ******** ***** ***** ************* ******

Bna_acc-C       CCTTATTAGTGAGAATGAGGAA---CAAGCTCAAGGAAGAGTGGACAAAGTTCTCAAAGA
Bna_acc-A       CCTTATTAGTGAGAATGAGGAGGACCAAGCTCAAGAAAGAGTGGAGAAAATTCTCAAAGA
                *********************    ********** ********* *** **********

Bna_acc-C       GGAAGAAGTAAGTTCGAGACTCCGTTCTGCAGGTGTCGGTGTGGTAAGCTGTATAATCCA
Bna_acc-A       GGAAGAAGTTAGTTCGAGCCTGTGTTCCGCAGGTGTGGGTGTGGTGAGCTGTATAATCGA
                ********* ******** **  **** ******** ******** ************ *
                                                                Q_ACC-A_F

Bna_acc-C       GCGGGATGAAGAACAAACGCCTATTAGACATTCGTTCCATTGGTCGATGGAGAAACAGTA
Bna_acc-A       GCGAGATGAAGGACGAACACCTATTAGACATTCGTTCCATTGGTCGATGGAGAAACAGTA
                *** ******* ** *** *****************************************
                                   Q_ACC-A_probe

Bna_acc-C       TTACGCAGAAGAGCCGATGCTGCGCCATCTTGAACCTCCTCTTTCCATATACCTTGAGTT
Bna_acc-A       CTATGCAGAAGAGCCGATGCTGCGCCATCTTGAACCTCCTCTCTCCATATACCTTGAGTT
                 ** ************************************** *****************
                     Q_ACC-A_R

Bna_acc-C       GGTATGAATGA-AACGTTAATAA--AATTCTTGTTTGACAAGCATTCTATATTCATGATC
Bna_acc-A       GGTAATGATAATAACGATCATCATGAAAACGTTACTAAAATGCTTGTTTATTTCATGATC
                ****   ** * **** * ** *  **  * *   * * * ** *  *   *********

Bna_acc-C       GTTTTTACTGTTTTGTCCGACGCAGGATAAGCTGAGAGGATATGAAAATATACAATATAC
Bna_acc-A       ATTTTTACTGTTTTGTCTGACGCAGGATAAGCTGAGAGGATATGAAAATATACAATATAC
                 **************** ******************************************

                          ACC_amp_R
Bna_acc-C       CCCTACAAGAGATCGTCAATGGCATCTG
Bna_acc-A       CCCTACAAGAGATCGTCAATGGCATCTG
                ****************************
```

**Supplementary Figure S4**. Sequence alignment of the two *ACC* gene copies in *B. napus* DH12075. Amplification primers for the 0.6 kb fragment are highlighted in green, whereas primers and probes specific for the *ACC*-A copy used in qPCR are highlighted in blue and yellow, respectively.

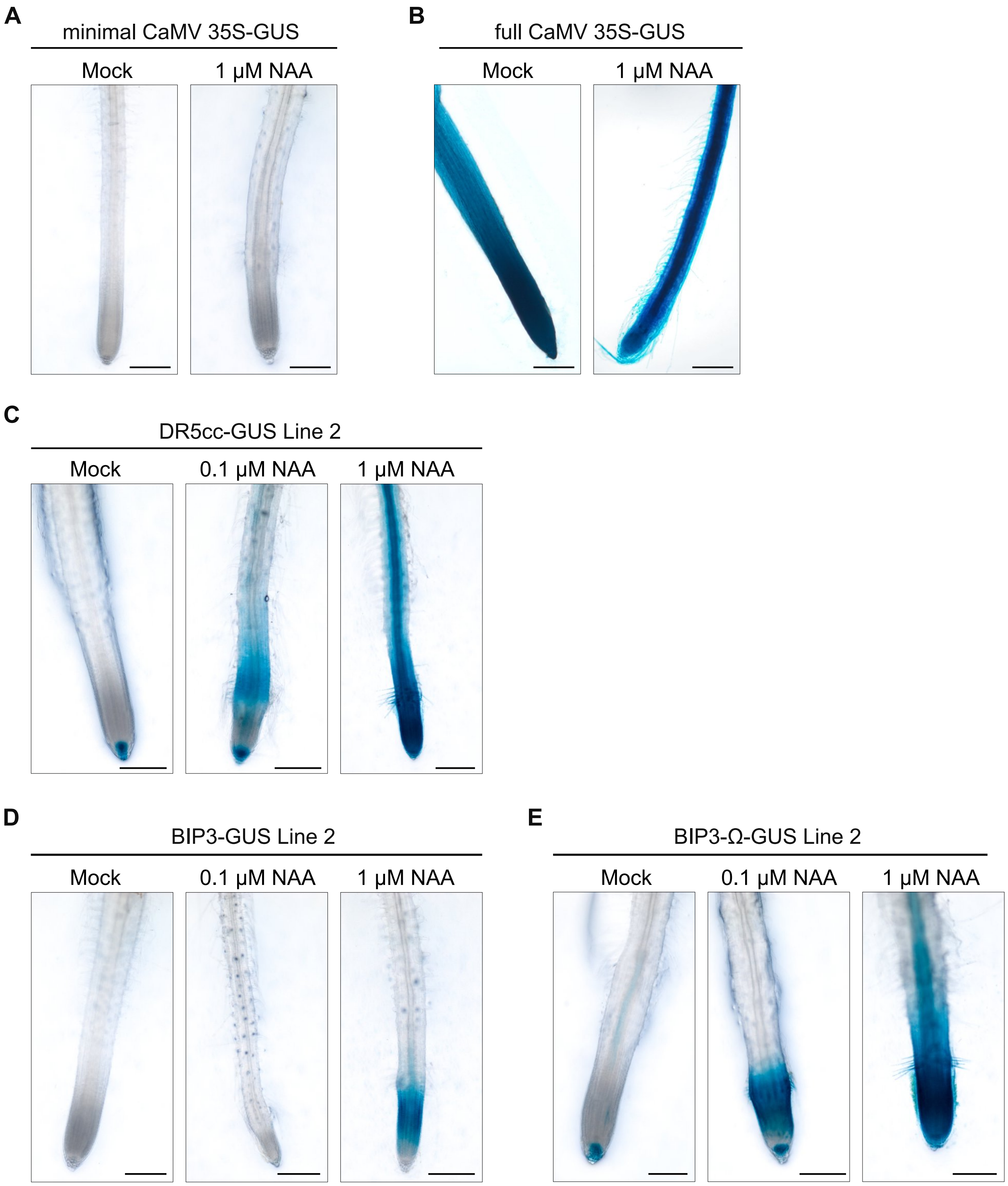


**Supplementary Figure S5**. GUS expression in hairy roots carrying reporter constructs driven by different promoters. Hairy roots expressing (**A**) the minimal CaMV 35S promoter-GUS reporter, (**B**) the full CaMV 35S promoter-GUS reporter, (**C**) the DR5cc-GUS reporter, Line 2 (three transgene copies), (**D**) BIP3-GUS, Line 2 (five transgene copies), and (**E**) BIP3-Ω-GUS, Line 2 (two transgene copies). Hairy roots were either mock-treated or treated with 0.1 μM or 1 μM NAA for 12 h prior to GUS staining. Scale bars represent 500 μm.

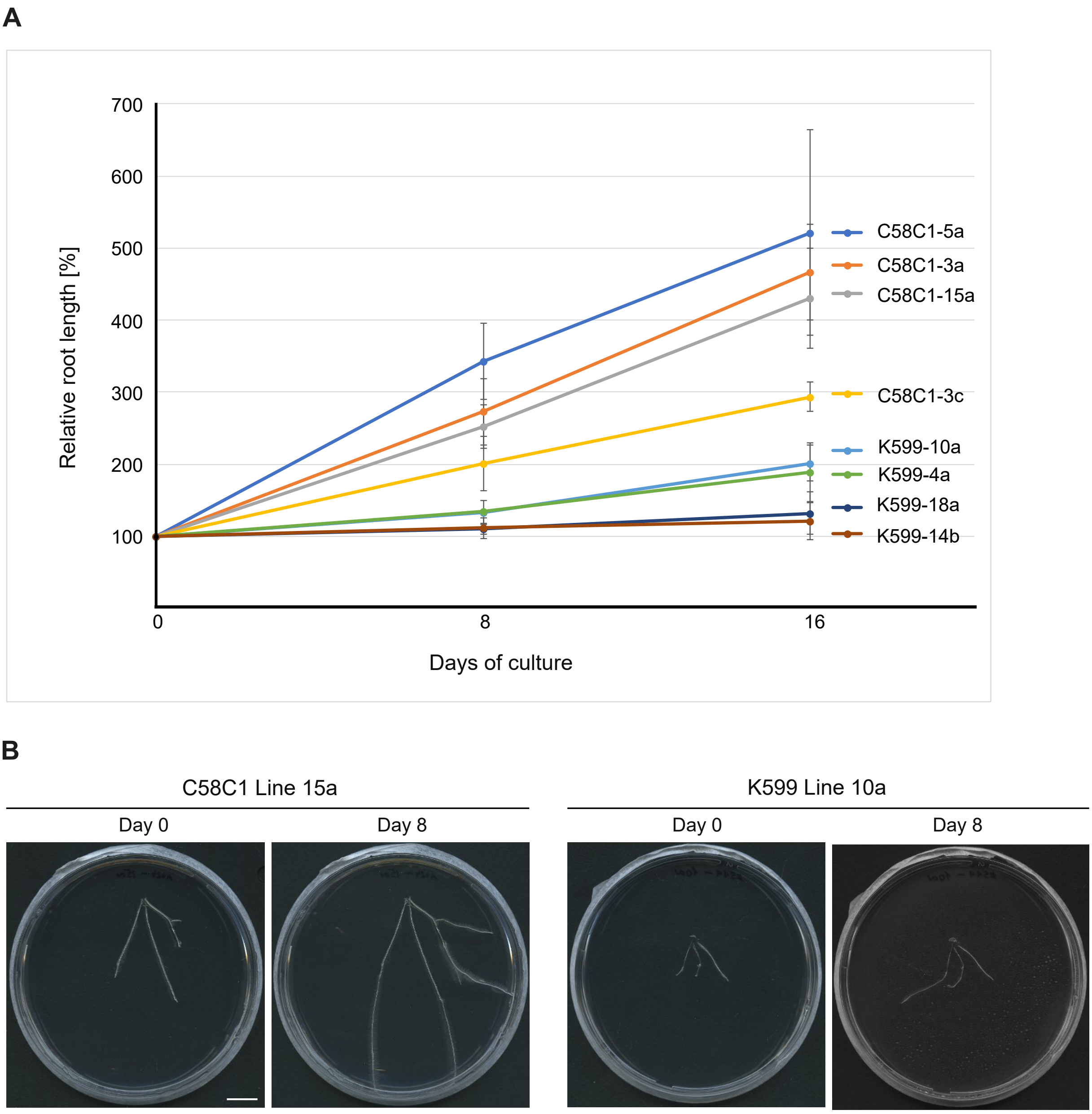


**Supplementary Figure S6**. Growth of hairy roots induced by the C58C1 or K599 *Agrobacterium* strains. (**A**) Relative root length (% of the initial root length at day 0) was measured after 8 and 16 days of culture in four lines transformed by C58C1 and four lines transformed by K599. Three independent biological replicates were performed, and the data are presented as mean values with standard deviation. (**B**) Representative images of hairy root lines at the initial time point and after 8 days of culture. Scale bar represents 1 mm.

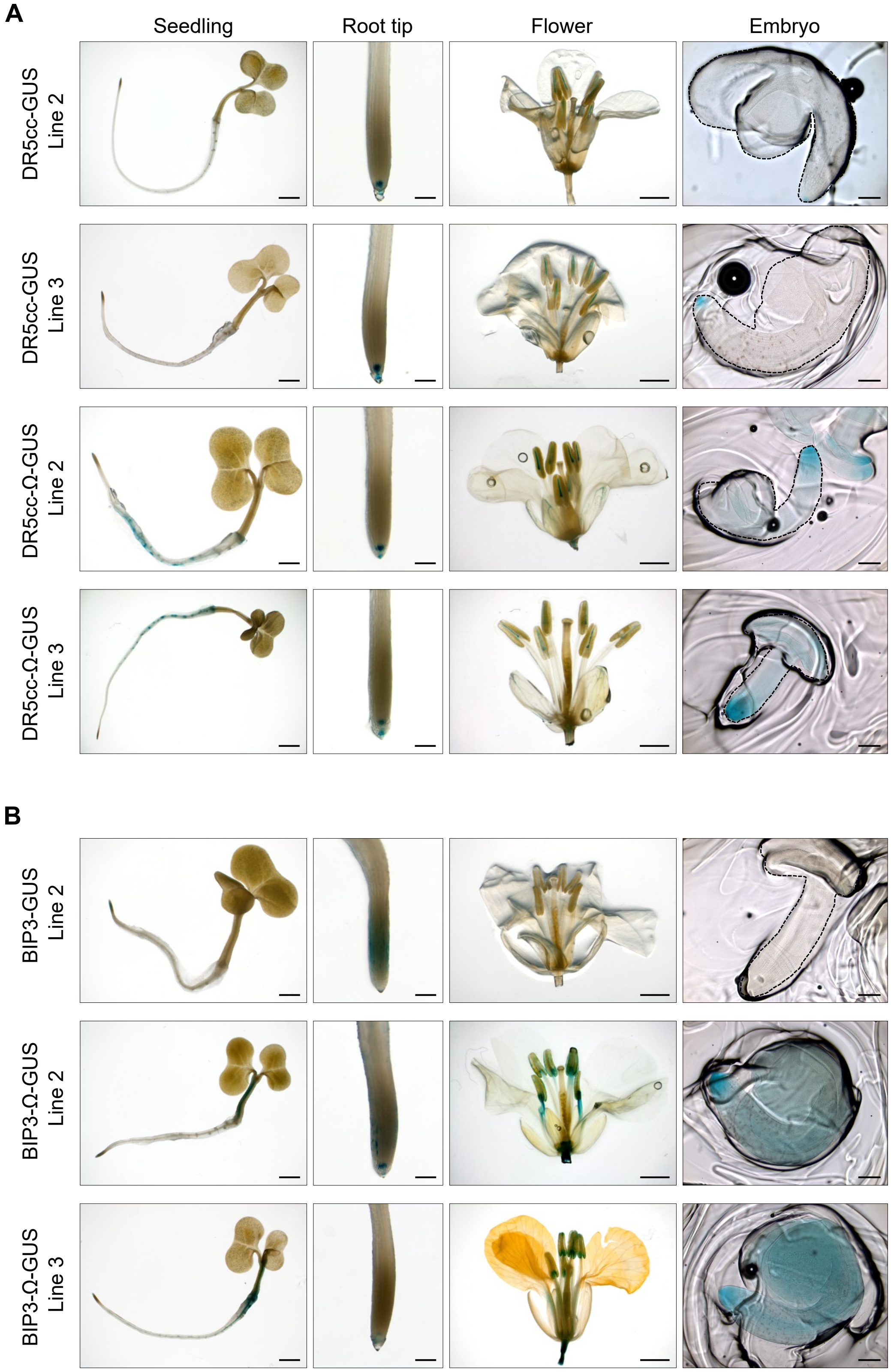


**Supplementary Figure S7**. Histochemical GUS staining of different tissues of *B. napus* transgenic plants. Independent transgenic lines from (**A**) DR5cc-GUS (+/- Ω) and (**B**) BIP3-GUS (+/- Ω) are presented. Scale bars represent 2 mm (seedlings, flowers), 200 μm (root tips), and 500 μm (embryos).

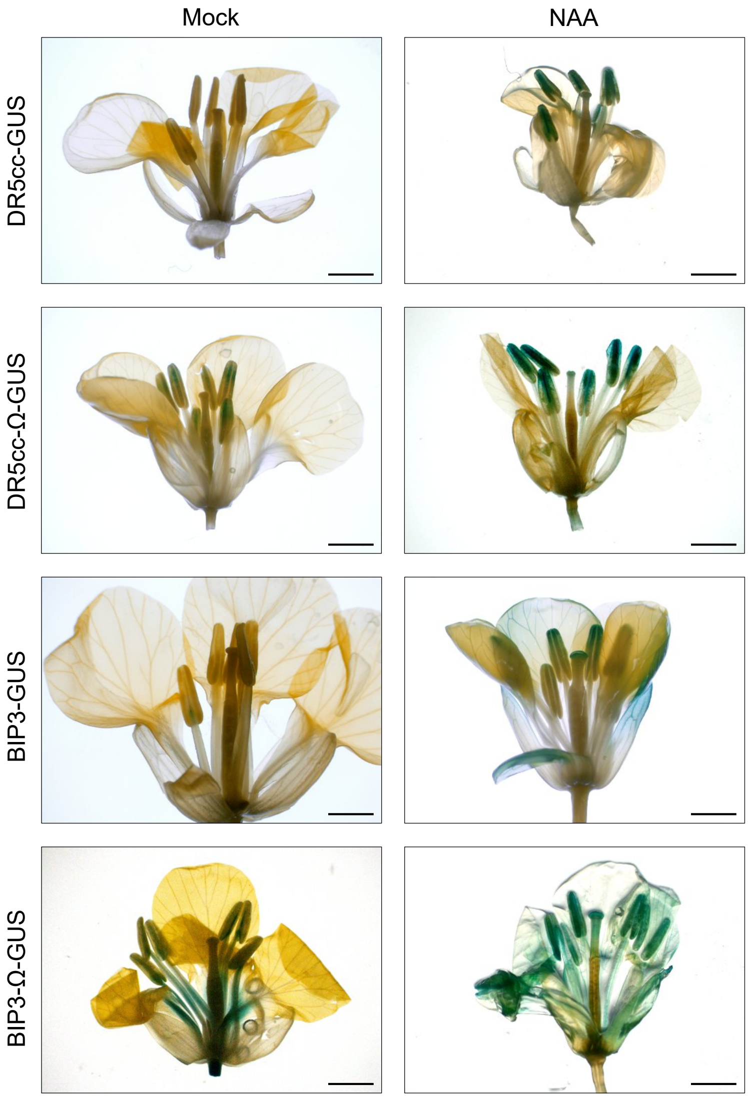


**Supplementary Figure S8**. Histochemical GUS staining of *B. napus* flowers expressing auxin reporters after NAA treatment. Flowers were sprayed with water (Mock) or 50 µM NAA in water, and stained after 24 h. Scale bars represent 2 mm.

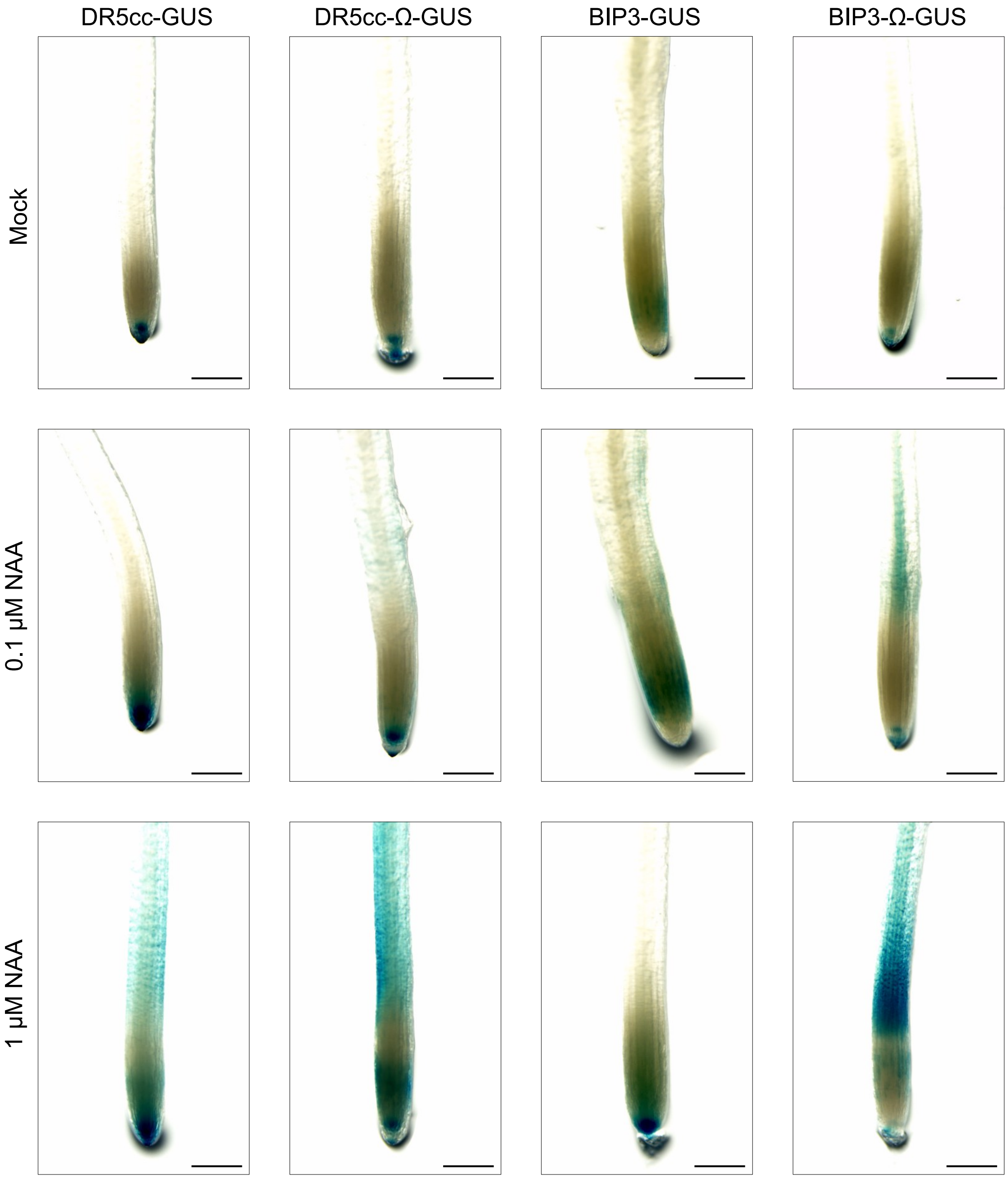


**Supplementary Figure S9**. Histochemical GUS staining of roots from regenerated transgenic *B. napus* plants following NAA treatment. Representative images showing GUS activity in root tips expressing DR5cc-GUS (+/- Ω) and BIP3-GUS (+/- Ω) under mock control conditions or following treatment with 0.1 μM and 1 μM NAA for 3 h. Scale bars represent 200 μm.

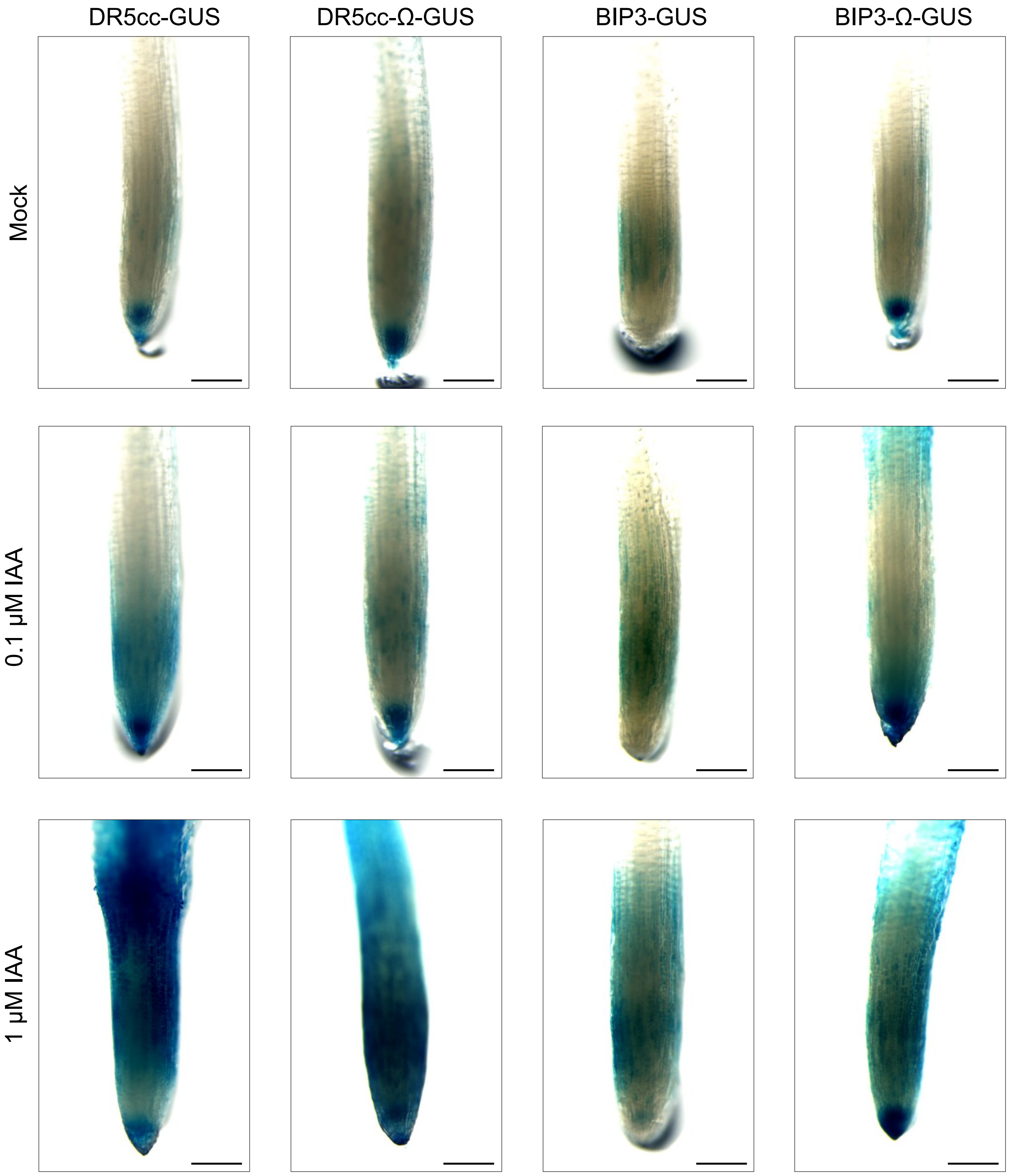


**Supplementary Figure S10**. Histochemical GUS staining of roots from regenerated transgenic *B. napus* plants following IAA treatment. Representative images showing GUS activity in root tips expressing DR5cc-GUS (+/- Ω) and BIP3-GUS (+/- Ω) under mock control conditions or following treatment with 0.1 µM and 1 µM IAA for 3 h. Scale bars represent 200 µm.

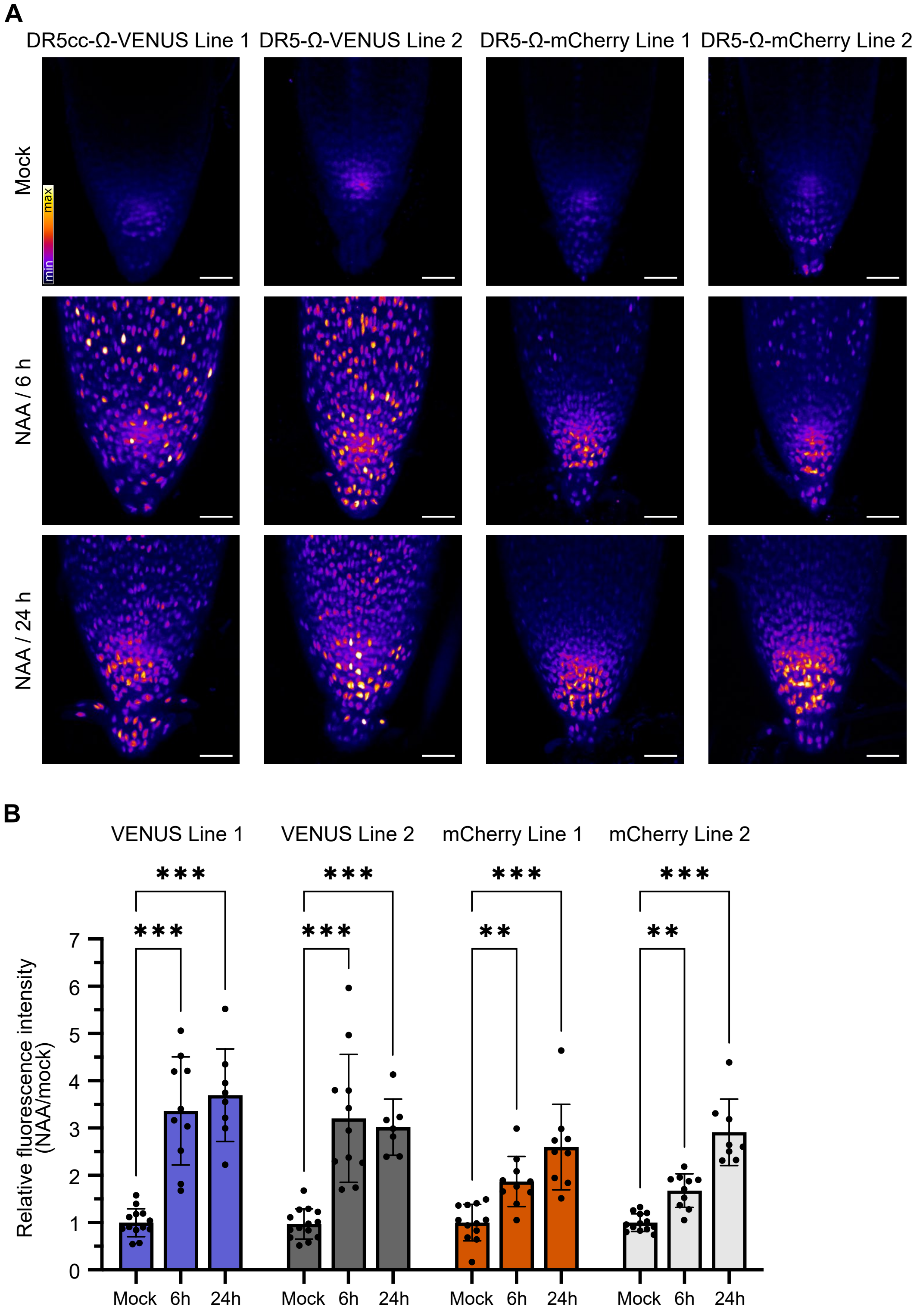


**Supplementary Figure S11**. The DR5cc reporter is responsive to NAA treatment in *B. napus* hairy roots. (**A**) Maximum intensity projections of root tips from independent lines expressing DR5cc-Ω-VENUS or DR5cc-Ω-mCherry treated with mock or 1 μM NAA, and imaged after 6 or 24 h. Scale bars represent 40 μm. (**B**) Quantification of relative VENUS and mCherry fluorescence intensities in mock- and NAA-treated hairy roots at indicated time points. Error bars indicate SD (n = 7-14 roots). For each time point, values are normalized to the corresponding mock. Statistical comparisons (one-way ANOVA with Dunnett's post-hoc test) were performed independently within each time point to compare stress treatments against the corresponding mock (**$P < 0.01$, ***$P < 0.001$).

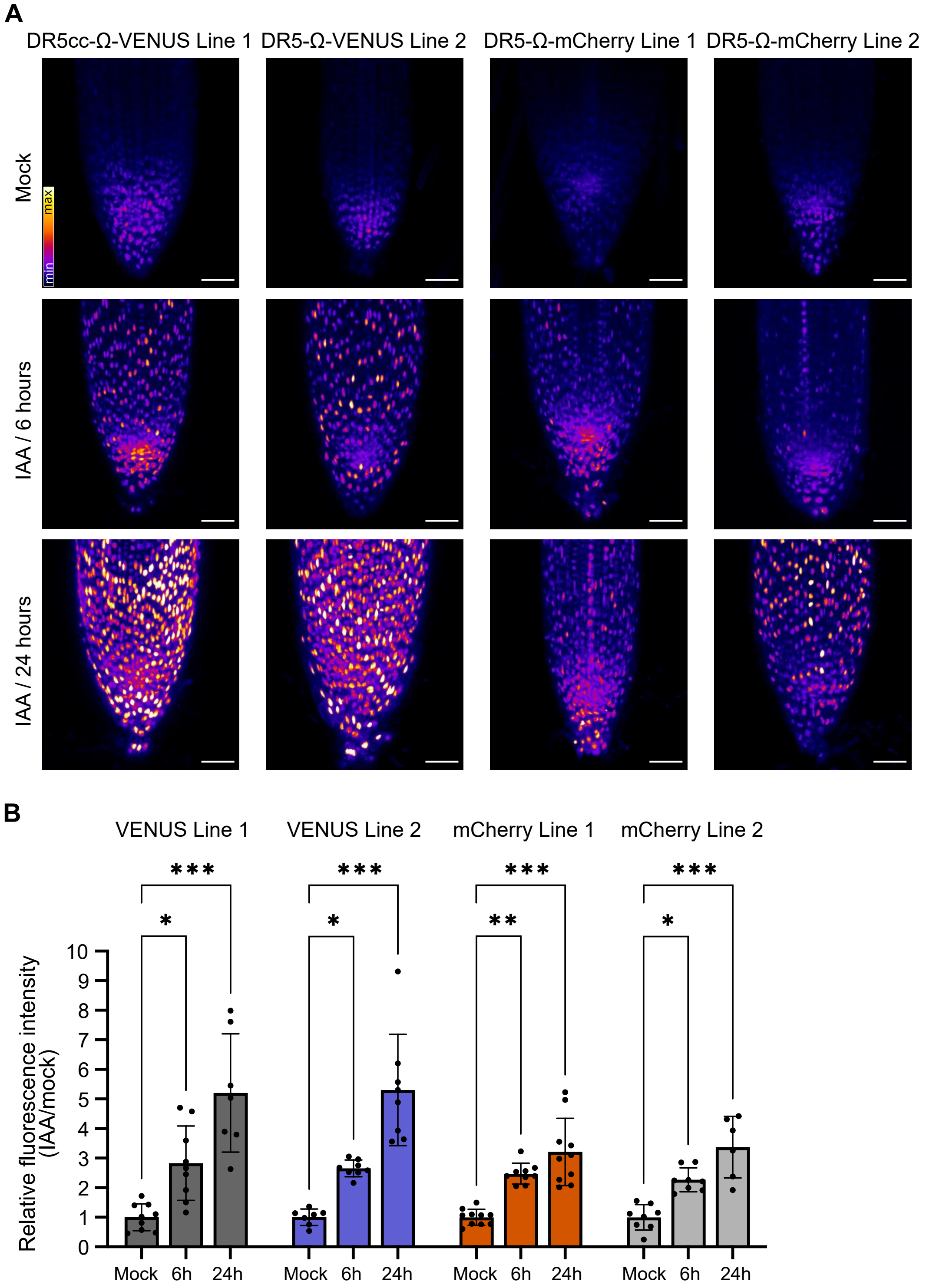


**Supplementary Figure S12**. The DR5cc reporter is responsive to IAA treatment in *B. napus* hairy roots. (**A**) Maximum intensity projections of root tips from independent lines expressing DR5cc-Ω-VENUS or DR5cc-Ω-mCherry treated with mock or 1 μM IAA, and imaged after 6 or 24 h. Scale bars represent 40 μm. (**B**) Quantification of relative VENUS and mCherry fluorescence intensities in mock- and IAA-treated hairy roots at indicated time points. Error bars indicate SD (n = 6-10 roots). For each time point, values are normalized to the corresponding mock. Statistical comparisons (one-way ANOVA with Dunnett's post-hoc test) were performed independently within each time point to compare stress treatments against the corresponding mock (*P < 0.5, **P < 0.01, ***P < 0.001).

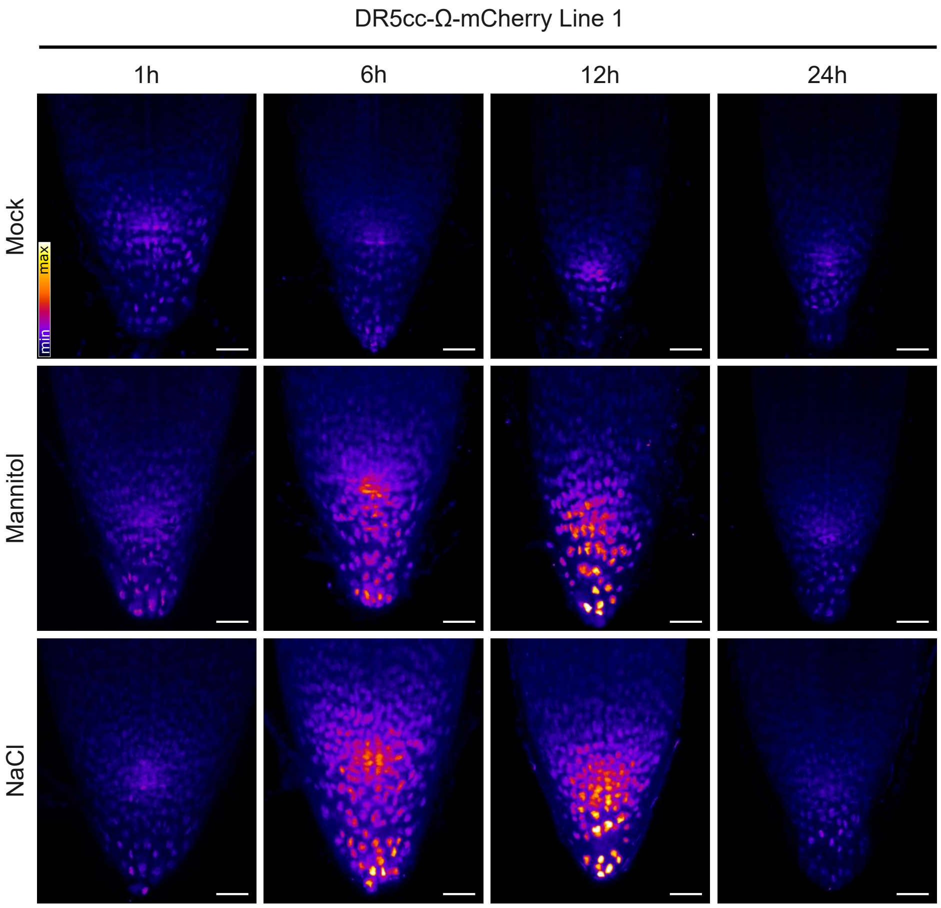


**Supplementary Figure S13**. Stress-induced auxin signaling dynamics in hairy roots expressing DR5cc-Ω-mCherry (Line 1). Maximum intensity projections of root tips treated with mock and abiotic stress (150 mM mannitol or 75 mM NaCl), and imaged at 1, 6, 12, and 24 h. Scale bars represent 40 μm. Quantification is shown in Figure 4.

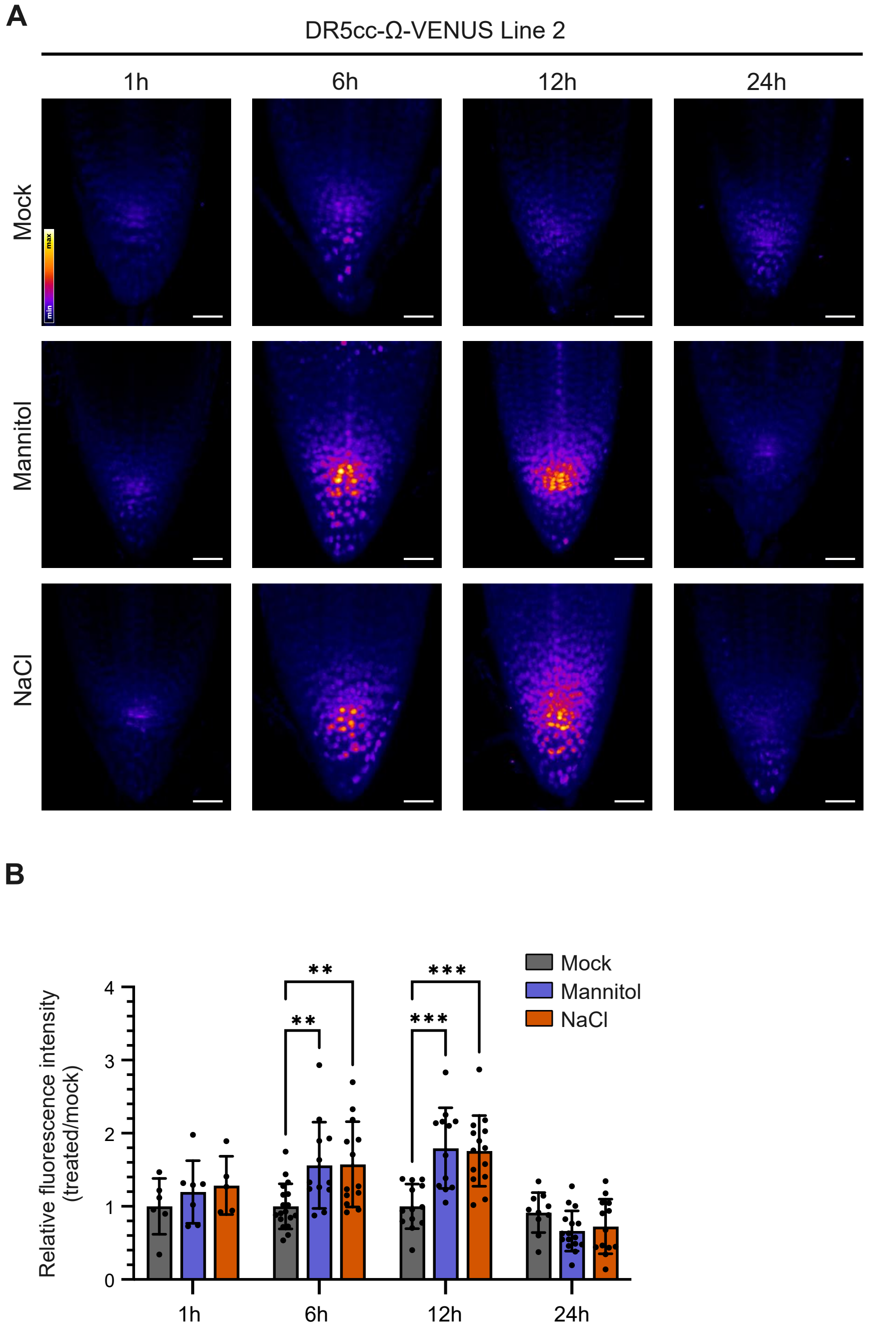


**Supplementary Figure S14**. Stress-induced auxin signaling dynamics in hairy roots expressing DR5cc-Ω-VENUS (Line 2). (**A**) Maximum intensity projections of root tips treated with mock and abiotic stress (150 mM mannitol or 75 mM NaCl), and imaged at 1, 6, 12, and 24 h. Scale bars represent 40 μm. (**B**) Quantification of relative VENUS fluorescence intensity in mock- and stress-treated hairy roots at indicated time points. Error bars indicate SD (n = 6-18 roots). For each time point, values are normalized to the corresponding mock. Statistical comparisons (one-way ANOVA with Dunnett's post-hoc test) were performed independently within each time point to compare stress treatments against the corresponding mock (**$P < 0.01$, ***$P < 0.001$); unannotated comparisons within a time point indicate no statistically significant difference ($P \geq 0.05$).

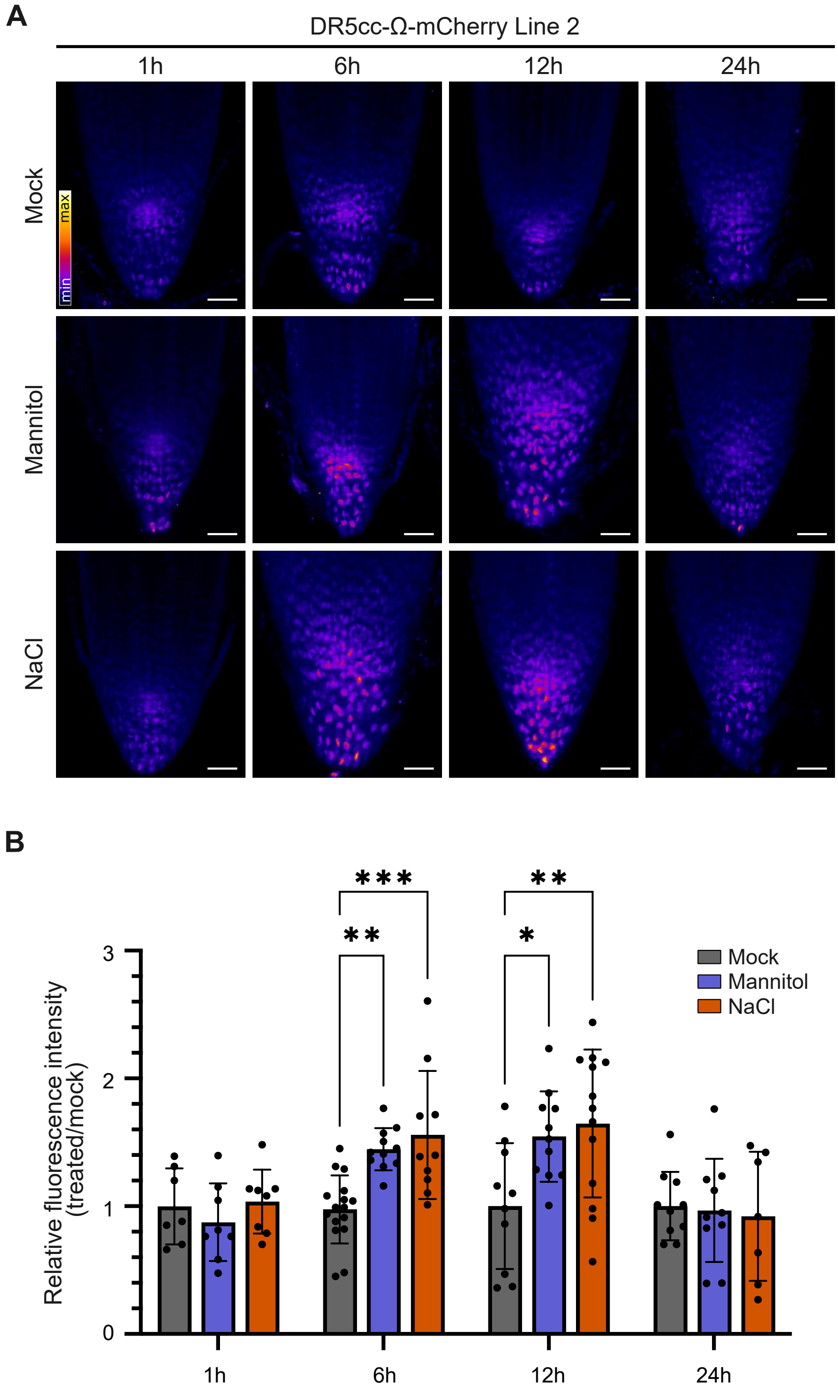


**Supplementary Figure S15**. Stress-induced auxin signaling dynamics in hairy roots expressing DR5cc-Ω-mCherry (Line 2). (**A**) Maximum intensity projections of root tips treated with mock and abiotic stress (150 mM mannitol or 75 mM NaCl), and imaged at 1, 6, 12, and 24 h. Scale bars represent 40 µm. (**B**) Quantification of relative mCherry fluorescence intensity in mock- and stress-treated hairy roots at indicated time points. Error bars indicate SD ($n = 7$-$16$ roots). For each time point, values are normalized to the corresponding mock. Statistical comparisons (one-way ANOVA with Dunnett's post-hoc test) were performed independently within each time point to compare stress treatments against the corresponding mock (*$P < 0.05$; **$P < 0.01$, ***$P < 0.001$); unannotated comparisons within a time point indicate no statistically significant difference ($P \geq 0.05$).

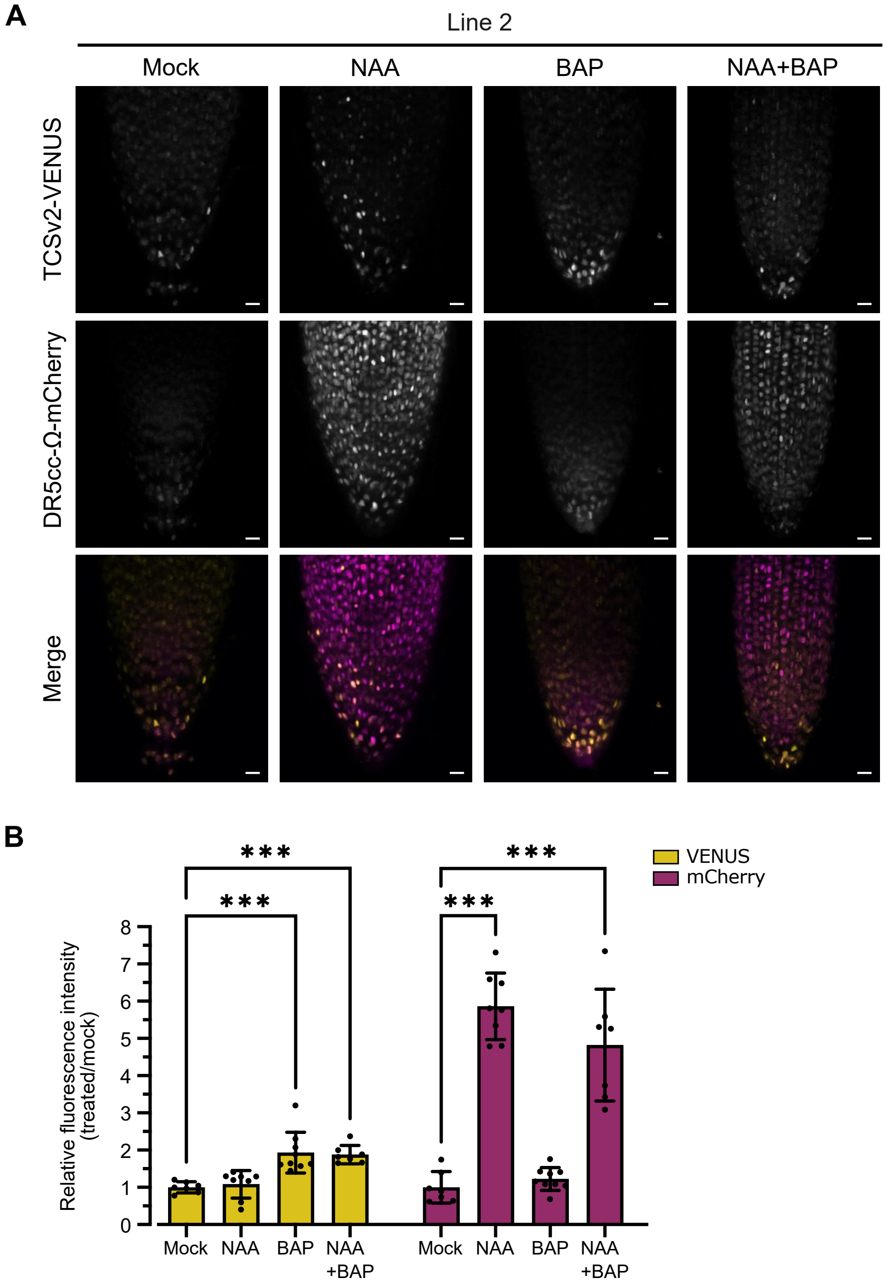


**Supplementary Figure S16**. Cytokinin and auxin reporter activity in *B. napus* hairy roots, transgenic Line 2. (**A**) Maximum intensity projections of hairy roots co-expressing TCSv2-VENUS and DR5cc-Ω-mCherry treated with mock, NAA (1 μM, 24 h), and/or BAP (5 μM, 2 h). Images are shown as VENUS (top row), mCherry (middle row) and merged channels (bottom row: magenta color for DR5cc-Ω-mCherry and yellow for TCSv2-VENUS). Scale bars represent 20 μm. (**B**) Quantification of relative VENUS (left) and mCherry (right) fluorescence intensities in mock- and hormone-treated hairy roots. Error bars indicate SD (n = 7-9 roots). For each channel, values are normalized to the corresponding mock. Statistical comparisons (one-way ANOVA with Dunnett's post-hoc test) were performed independently within each channel to compare hormonal treatments against the corresponding mock (***$P < 0.001$); unannotated comparisons within a time point indicate no statistically significant difference ($P \geq 0.05$).

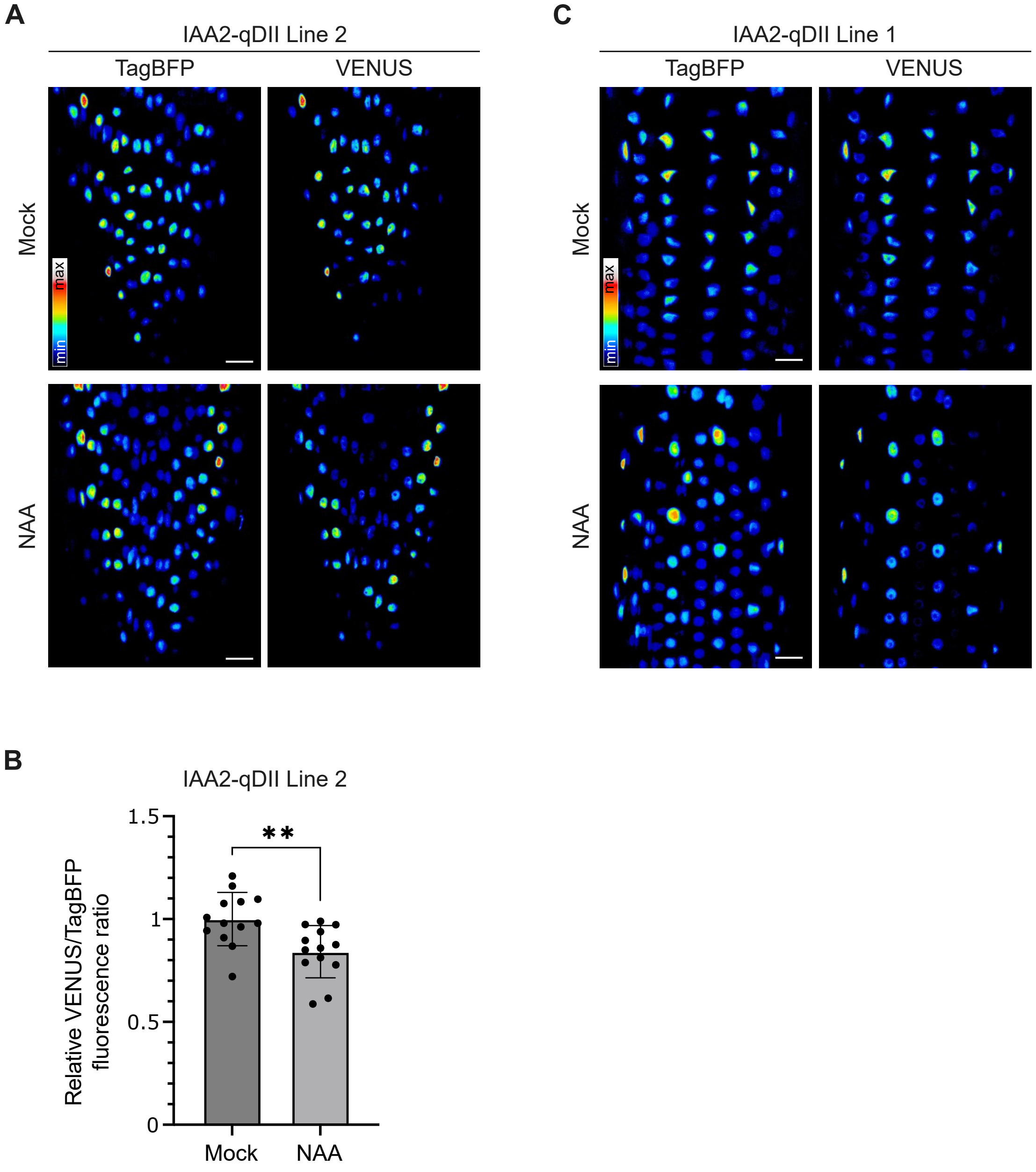


**Supplementary Figure S17**. Auxin responsiveness of the IAA2-qDII reporter in *B. napus* hairy roots following NAA treatment. (**A**) Representative images of epidermal cells from IAA2-qDII Line 2 under mock and NAA (10 μM, 2 h) conditions, showing TagBFP (left) and VENUS (right) fluorescence. (**B**) Quantification of the nuclear VENUS/TagBFP fluorescence intensity ratio for the conditions shown in (A) (**P < 0.01, Welch's t-test). Values are normalized to the corresponding mock. Error bars indicate standard deviation (SD; n = 13 roots per treatment). (**C**) Representative images of epidermal cells from IAA2-qDII Line 1 under mock and NAA (10 μM, 2 h) conditions, showing TagBFP (left) and VENUS (right) fluorescence. The corresponding quantification of the nuclear VENUS/TagBFP fluorescence intensity ratio is presented in Figure 6D. Scale bars represent 20 μm.

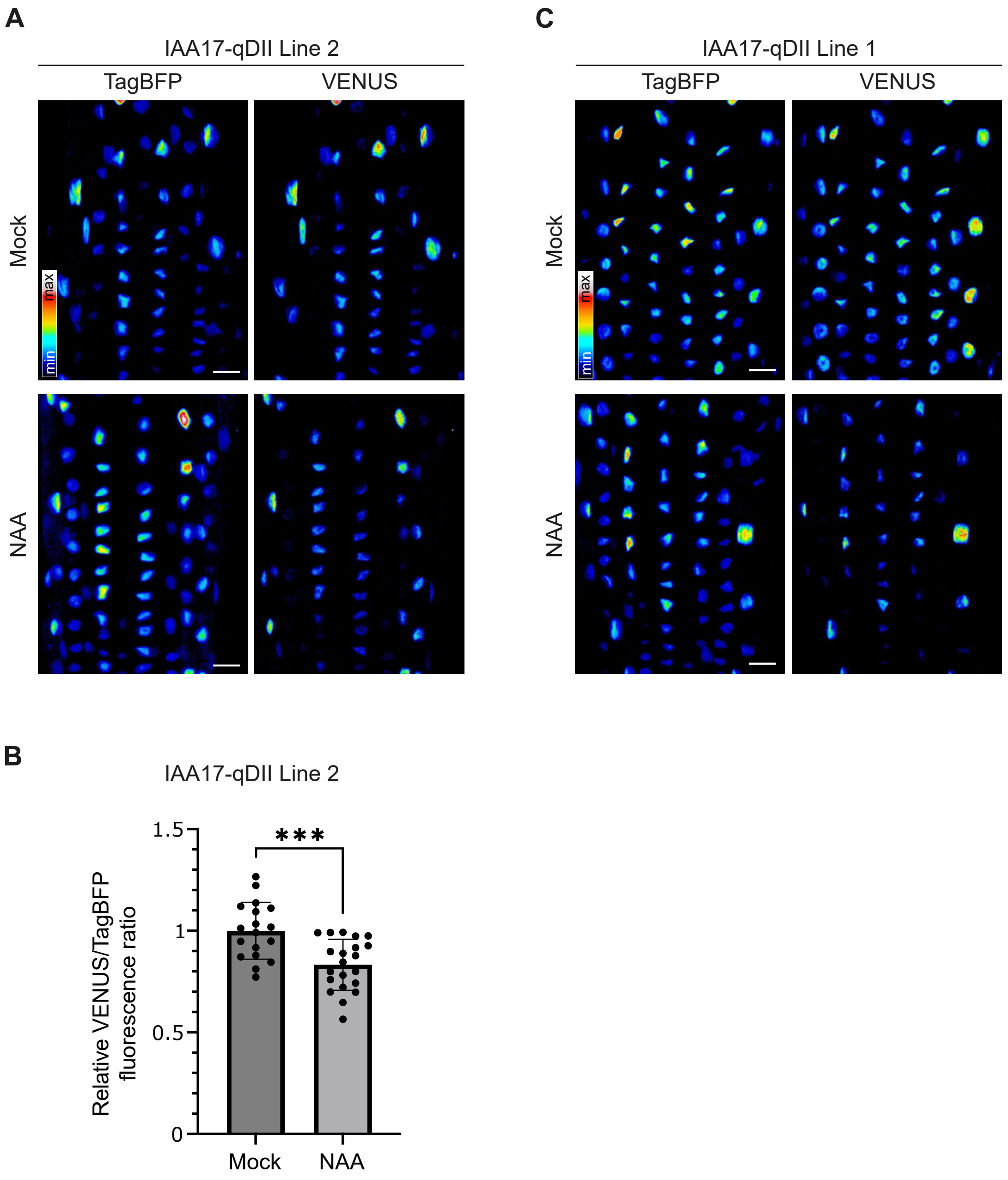


**Supplementary Figure S18**. Auxin responsiveness of the IAA17-qDII reporter in *B. napus* hairy roots following NAA treatment. (**A**) Representative images of epidermal cells from IAA17-qDII Line 2 under mock and NAA (10 μM, 2 h) conditions, showing TagBFP (left) and VENUS (right) fluorescence. (**B**) Quantification of the nuclear VENUS/TagBFP fluorescence intensity ratio for the conditions shown in (A) (**P < 0.01, Welch's t-test). Values are normalized to the corresponding mock. Error bars indicate standard deviation (SD; n = 18 − 21 roots). (**C**) Representative images of epidermal cells from IAA17-qDII Line 1 under mock and NAA (10 μM, 2 h) conditions, showing TagBFP (left) and VENUS (right) fluorescence. The corresponding quantification of the nuclear VENUS/TagBFP fluorescence intensity ratio is presented in Figure 6F. Scale bars represent 20 μm.

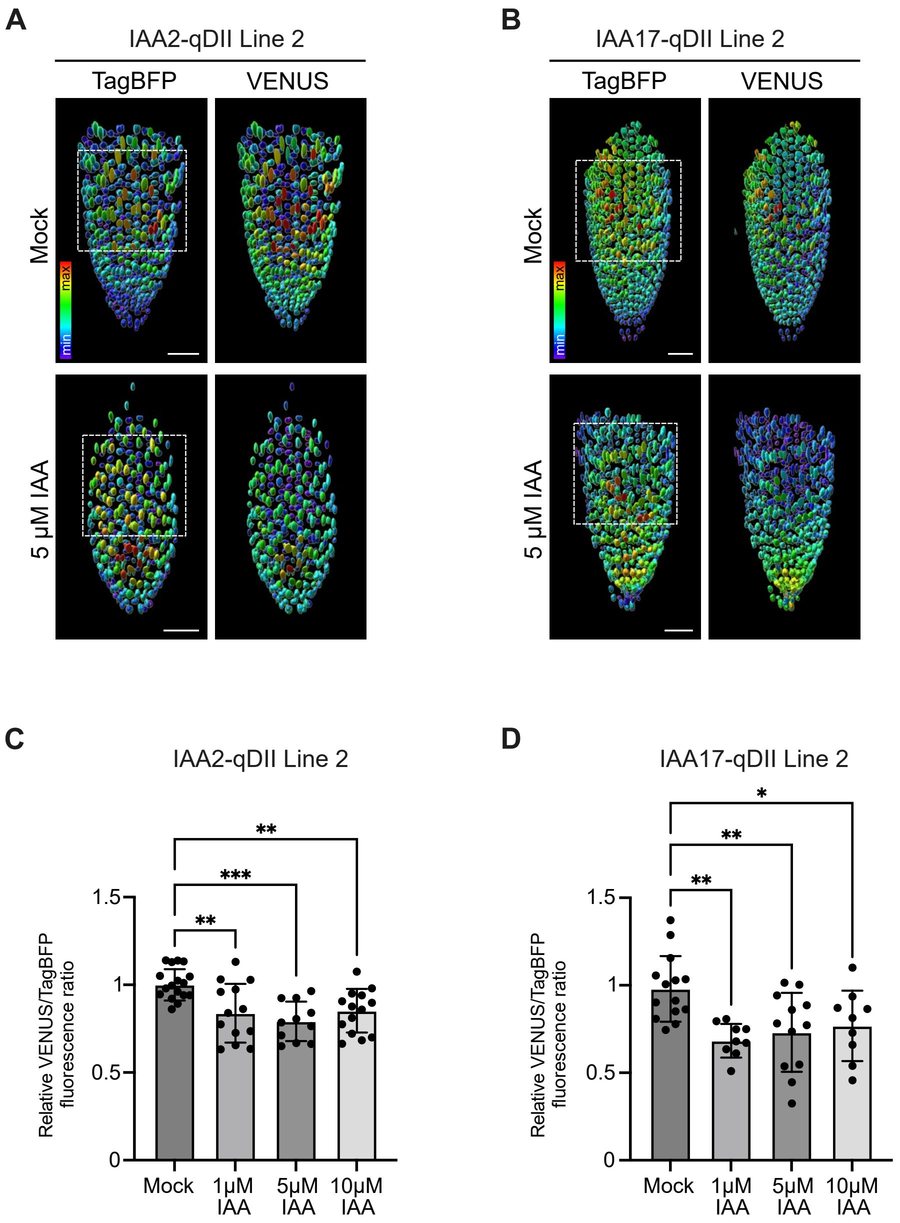


**Supplementary Figure S19**. IAA responsiveness of IAA2-qDII Line 2 and IAA17-qDII Line 2 in *B. napus* hairy roots. Maximum intensity projections of hairy root tips from (**A**) IAA2-qDII Line 2 and (**B**) IAA17-qDII Line 2 under mock and IAA (5 µM, 2 h) treatment conditions, showing measured signal intensities of TagBFP (left) and VENUS (right). The boxed region indicates the area used for nuclear signal quantification. Images were generated using IMARIS software. Scale bars represent 50 µm. Quantification of the nuclear VENUS/TagBFP fluorescence intensity ratio for (**C**) IAA2-qDII Line 2 and (**D**) IAA17-qDII Line 2. Statistical significance was assessed by one-way ANOVA followed by Dunnett's post hoc test (*$P < 0.05$, **$P < 0.01$, ***$P < 0.001$). Values are normalized to the corresponding mock controls. Error bars indicate standard deviation (SD; n = 9 – 17 roots).

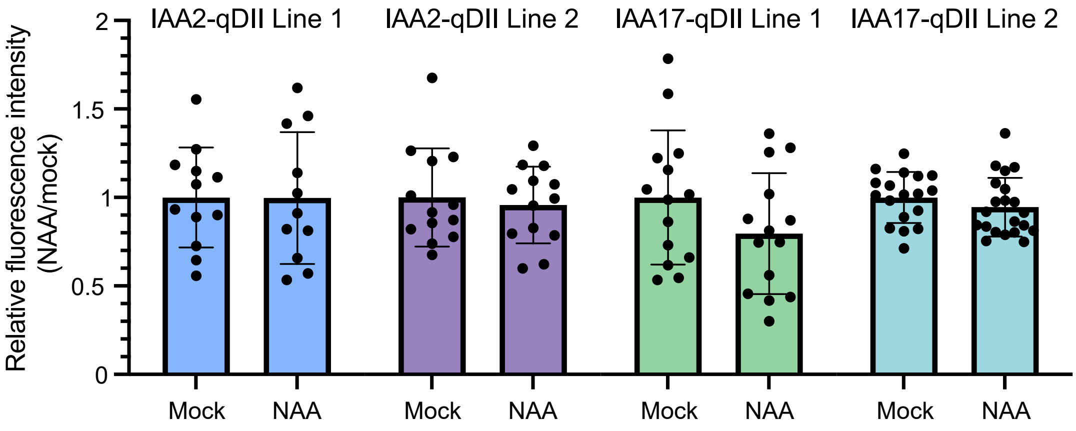


**Supplementary Figure S20**. Stability of the TagBFP fluorescence in qDII constructs. Quantification of relative TagBFP fluorescence intensity in independent hairy root lines expressing IAA2 / IAA17-qDII treated with mock or NAA (10 μM, 2 h). Error bars indicate SD (n = 11 − 21 roots). No significant differences were observed in treated samples compared with the mock control (Welch's t-test).

**Supplementary Table S1.** List of oligonucleotides.

**Genotyping of T1 plants**

| | |
|---|---|
| GUS_F | TGCTCTACACCACGCCGAACA |
| GUS_R | GAGCATCTCTTCAGCGTAAGGG |
| TL_rolA_F | GTTAGGCGTGCAAAGGCCAAG |
| TL_rolA_R | TGCGTATTAATCCCGTAGGTC |
| TR_aux1_F | CATAGGATCGCCTCACAGGT |
| TR_aux1_R | CGTTGCTTGATGTCAGGAGA |

**Transgene copy number assessment**

| | |
|---|---|
| ACC_amp_F | ATCGCTGGAGTTTCTGCCGAA |
| ACC_amp_R | CAGATGCCATTGACGATCTCTTGTAG |
| Q_ACC-A_F | GTGAGCTGTATAATCGAGCGA |
| Q_ACC-A_R | GCATCGGCTCTTCTGCATAG |
| Q_ACC-A_probe | AGGACGAACACCTATTAGACATTCGTTCCATT |
| Q_GUS_F | GACGAAAACGGCAAGAAAAAGCAG |
| Q_GUS_R | TGAAGTACCTGCCAGTCAACAG |
| Q_GUS_probe | AATGCTCTACACCACGCCGAACACCTG |
